# Cell-sized confinement controls generation and stability of a protein wave for spatiotemporal regulation in cells

**DOI:** 10.1101/641662

**Authors:** Shunshi Kohyama, Natsuhiko Yoshinaga, Miho Yanagisawa, Kei Fujiwara, Nobuhide Doi

## Abstract

Min system, which determines the cell division plane of bacteria, uses the localization change of protein (Min wave) emerged by a reaction-diffusion coupling. Although previous studies have shown that cell-sized space and boundaries modulate shape and speed of Min waves, its effects on Min wave emergence was still elusive. Here, by using a fully confined microsized space as a mimic of live cells, we revealed that confinement changes conditions for Min wave emergence. In the microsized space, an increase of surface-to-volume ratio changed the localization efficiency of proteins on membranes, and therefore, suppression of the localization change was necessary to produce stable Min wave generations. Furthermore, we showed that the cell-sized space more strictly limits parameters for wave emergence because confinement inhibits instability and excitability of the system. These results illuminate that confinement of reaction-diffusion systems works as a controller of spatiotemporal patterns in live cells.

## Introduction

Spatiotemporal self-organization of biomolecules in cells takes on a fundamental mechanism to maintain cellular structure. In particular, the intracellular reaction-diffusion wave (iRD) is an essential mechanism for various spatiotemporal regulation including DNA segregation (1), cell-shape deformation, cell migration (2, 3), and cell polarization (4). A remarkable example of iRD is Min wave, which is a bacterial spatiotemporal organization system (Min system). The Min system precisely places the division site at the center of the cell by using iRD (5, 6). This system comprises three proteins called MinC, MinD, and MinE with localization of MinD and MinE oscillating between one pole to the other by a coupling between biochemical reactions and molecular diffusions (6, 7). MinC has no role in the Min wave but rather inhibits polymerization of a cell division initiation factor (FtsZ) by following the Min wave. This process enforces initiation of cell division only at the center of cells (5, 6).

To date, Min wave is the only biological RD system reconstituted *in vitro*. Reconstitution of Min wave was firstly shown by spotting a mixture of MinD, MinE, and ATP on 2-demensinal (2D) planar membranes comprising *E. coli* polar lipid extract in open geometry(7). The following studies based on a 2D planar system have clarified the mechanisms of wave generation and the characteristics of Min waves (8–13). *In vitro* studies have demonstrated that external environments such as boundary shape, protein concentration, and lipids alter patterns, velocities, wavelengths and shapes of Min waves (9–14). In particular, boundary shapes prepared by PDMS chambers significantly change the behavior of Min waves with studies showing that a rod-shape is important in terms of inducing the pole-to-pole oscillation found in living cells (10, 12, 13).

Due to the importance of Min waves for initiating division at a precise location, the timing, conditions, and regulation of their emergence should be investigated. However, despite the critical conditions for Min wave emergence including environmental effects having been surveyed in open space, the effect of confinement in cell-sized space, which is one of the most remarkable features of living cells, has been poorly addressed. Although some studies have reported reconstitution of Min waves in fully confined cell-sized spaces (12, 13, 15), lipid conditions were modified or spaces were closed after observing wave generation. The necessity of these treatments suggests that cell-sized space affects the condition for Min wave emergence.

Recent studies have unveiled that confinement inside cell-sized space alters both behaviors of biochemical reactions and molecular diffusions (16–18). Because RD waves appear only in limited parameter ranges (19, 20), encapsulation inside cell-sized space should shift the condition for Min wave emergence such as diffusion and interaction of its elements. Moreover, considering that interference of the RD waves at the time of two-wave collision (21) and initiation of Min waves by interaction among Min proteins on 2D planar membranes (7), it is plausible that the condition for wave emergence in a small space for only a single wave is different from that in large spaces for multiple waves.

In this study, we investigated the generation mechanism of Min waves in micro-sized closed space fully covered with *E. coli* polar lipid extract. Our experimental and theoretical analyses revealed that a fully confined micro-sized space changes the rate of protein localizations, and therefore, elements to cancel the effect were necessary to produce Min waves in a small space. Furthermore, our results show that cell-sized space itself enrolls on spatio-temporal regulation via RD mechanisms in living cells by making the emergence mechanism distinct from open system such as *in vitro*.

## Results

### MinDE are insufficient for emerging Min waves in micro-sized space fully covered with E. coli polar lipid extract

Previous works have reported that only MinD, MinE, and ATP are necessary and sufficient for generation of Min waves on 2D planar membranes of *E. coli* polar lipid extract (7) (Figure 1A) and in fully confined cell-sized space by using a modified lipid mixture(12, 15). The Min wave in open geometry has been well characterized in many laboratories (8, 11, 13), and has also been reproduced by using materials prepared in our laboratory (sfGFP-MinD and MinE-mCherry)(Figure 1B). Then, we encapsulated these materials in micro-sized space fully covered with *E. coli* polar lipid extract using an emulsification method (12, 22). However, we found that sfGFP-MinD, MinE-mCherry, and ATP are insufficient for Min wave emergence in the microdroplets covered with *E. coli* polar lipids (Figure 1C, Video 1). Usage of non-fluorescent tagged MinDE tracked by sfGFP-MinC indicated that fluorescent proteins fused with MinD or MinE could not explain the lack of wave occurrence (Figure 1—figure supplement 1AB). In contrast, sfGFP-MinD, MinE-mCherry, and ATP induced Min waves in microdroplets covered with a lipid mixture (85% DOPC and 15% Cardiolipin), as reported previously (12) (Video 2). These results indicated that encapsulation of micro-sized space fully covered with lipid alters some critical parameters for Min wave emergence compared to the case for 2D membranes.

**Figure 1:**
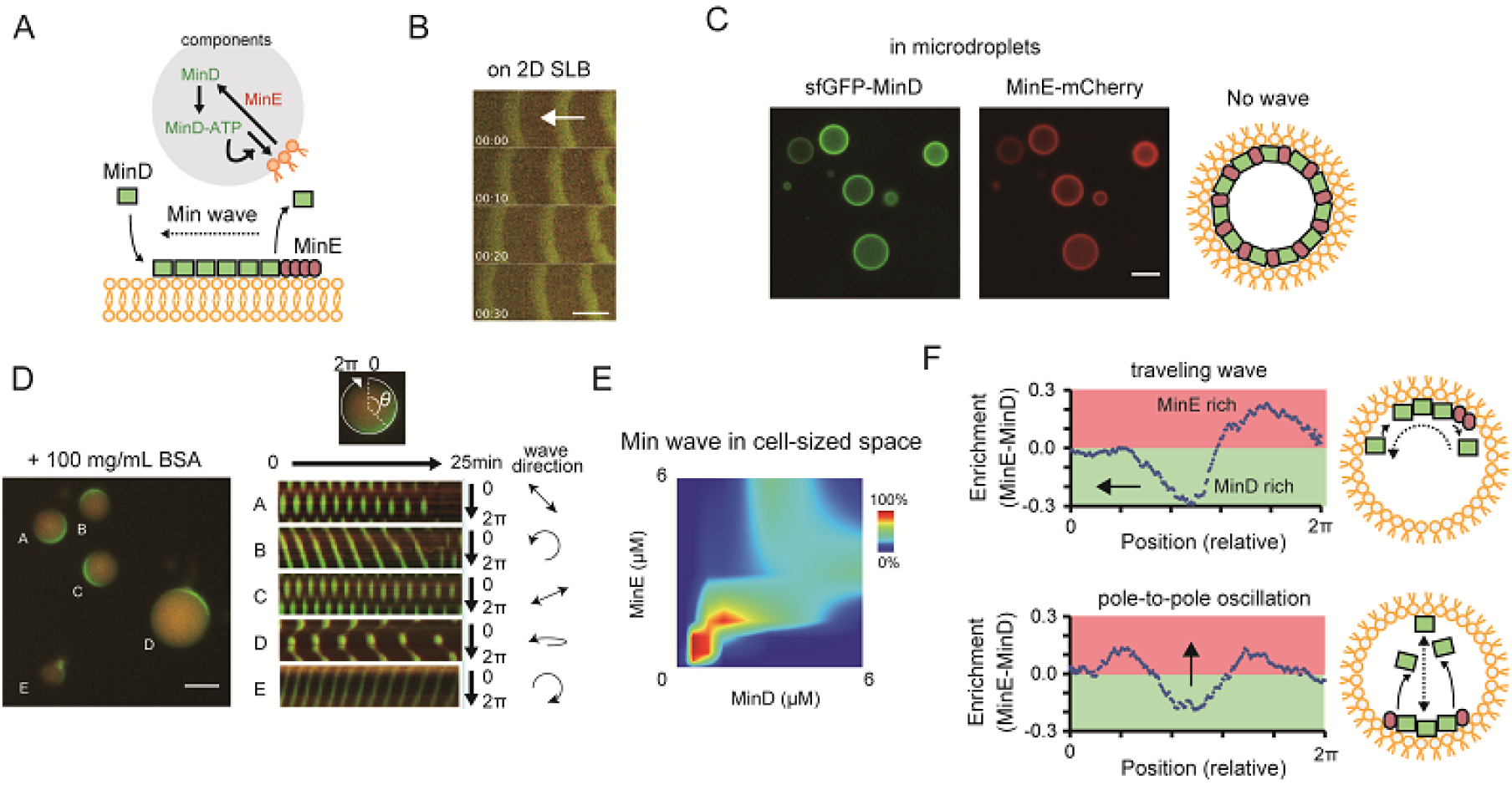
Min waves emergence in microdroplets as a result of high concentration BSA addition. (**A**) Schematic illustration of a simplified molecular mechanisms underlying Min wave propagation and the experimental system. (**B**) Min wave on 2D supported lipid bilayers (SLB). (**C,D**) Microdroplets encapsulating 1 μM sfGFP-MinD, MinE-mCherry, and 2.5 mM ATP in the absence (**C**) or the presence of 100 mg/mL BSA (**D**). Scale bars: 10 μm. Kymographs of sfGFP-MinD (green) and MinE-mCherry (red) in the proximity of membranes in each droplet shown at the right of each panel. Kymographs were generated by tracking fluorescence intensities along circumference lines on membrane surface. Arrows beside the kymographs show the direction and mode of Min wave. Single-round and double-headed arrows indicate traveling wave and pole-to-pole oscillation, respectively. (**E**) Probability of inhomogeneous localization and wave propagation revealed by the reconstitution experiments at various concentrations of sfGFP-MinD and MinE-mCherry in microdroplets. (**F**) Enrichment profiles of MinD and MinE derived from normalized surface plots.

### Addition of protein crowder assists Min wave emergence in cell-sized droplets

The difference of emergence conditions between being located on 2D and in 3D closed space raised a possibility that some factors other than MinDE regulates Min wave emergence in living cells. From the fact that lipid species change conditions for Min wave emergence, we assumed that changes of balance between reaction and diffusion by some factors such as physicochemical environments are associated with this difference.

As a candidate for such a factor, we focused on molecular crowding in cells. In living cells, ~30% of cell mass consists of macromolecules, and such crowding modulates both biochemical reactions and molecular diffusion (23, 24). Thereby, crowding is likely associated with patterns associated with reaction-diffusion systems. In fact, crowding agents that emulate molecular crowding *in vitro* have been shown to affect coupling of Min waves over membrane gaps (25) and wavelength of Min waves in the presence of FtsZ system (26).

To test this possibility, synthetic polymers (PEG8000 or Ficoll70), or a protein-based crowding agent (BSA) were mixed with MinDE and ATP with the mixtures then being encapsulated in microdroplets covered with *E. coli* polar lipid. Remarkably, co-supplementation of BSA at high concentration (100 mg/mL) with Min proteins induced Min wave emergence (Figure 1D, Video 3), while neither PEG8000 nor Ficoll70 induced Min waves (Video 4 and 5). Supplementation of BSA also induced Min wave emergence in the case of no fluorescence-tagged MinDE being tracked by sfGFP-MinC (Figure 1—figure supplement 1C, Video 6).

Varying concentrations of MinD and MinE indicated that both proteins should be around 1 μM to lead the emergence of Min waves (Figure 1E), consistent with their concentrations *in vivo*(27). ATP replacement with ADP or ATPγS, or replacement of MinD with an ATPase-deficient mutant(28), showed that the Min wave depends on ATP (Figure 1—figure supplement 2), similar to the case for 2D planar membranes (7). The frequent patterns observed were pole-to-pole oscillations and traveling waves (Figure 1—figure supplement 3), as noted by a previous study using modified lipids (12). MinE was enriched at the tail of the traveling wave (Figure 1F top) and was enriched at both tails of the wave in the case of pole-to-pole oscillations (Figure 1F bottom). These MinE enrichments were similar to the previous reports of traveling waves on 2D planar membranes (7), and the E ring observed in living cells with such pole-to-pole oscillations (5).

### BSA modifies attachment of MinE on membranes without MinD

To understand why the emergence condition for Min wave differs between being located on 2D planar membranes and in 3D closed geometry, we investigated the mechanism of wave emergence induced by BSA in micro-sized closed space. It has been known that crowding agents changes reaction rates, diffusion rates, and increase association rates by depletion force (23, 24). Thus, we first assumed that BSA may change reaction rates of MinD and MinE binding each other or to the membrane, decrease diffusion constants of MinD and MinE in the cytosol, and increase association rates between macromolecules by a depletion force. Among these effects, the effect of depletion force is excluded because previous studies have indicated that BSA shows little depletion force in contrast to PEG8000 or Ficoll70, which show a strong depletion force (24).

First, diffusion of macromolecules inside closed space was evaluated by using fluorescence correlation spectroscopy (FCS) and Fluorescence Recovery After Photo-bleaching (FRAP). FCS revealed that diffusion rates of GFP in cytosolic parts at 50 mg/mL BSA were similar to that in non-crowding conditions, but decreased at over 100 mg/mL (Figure 2A). However, we found that Min waves were stably generated even at 50 mg/mL of BSA (Figure 2—figure supplement 1). The effect of BSA on diffusion of sfGFP-MinD on membranes was investigated by FRAP and it was found that even 100 mg/mL of BSA slightly but did not significantly decrease the diffusion rates of MinD on lipid membranes of various sizes of microdroplets (Figure 2B).

**Figure 2:**
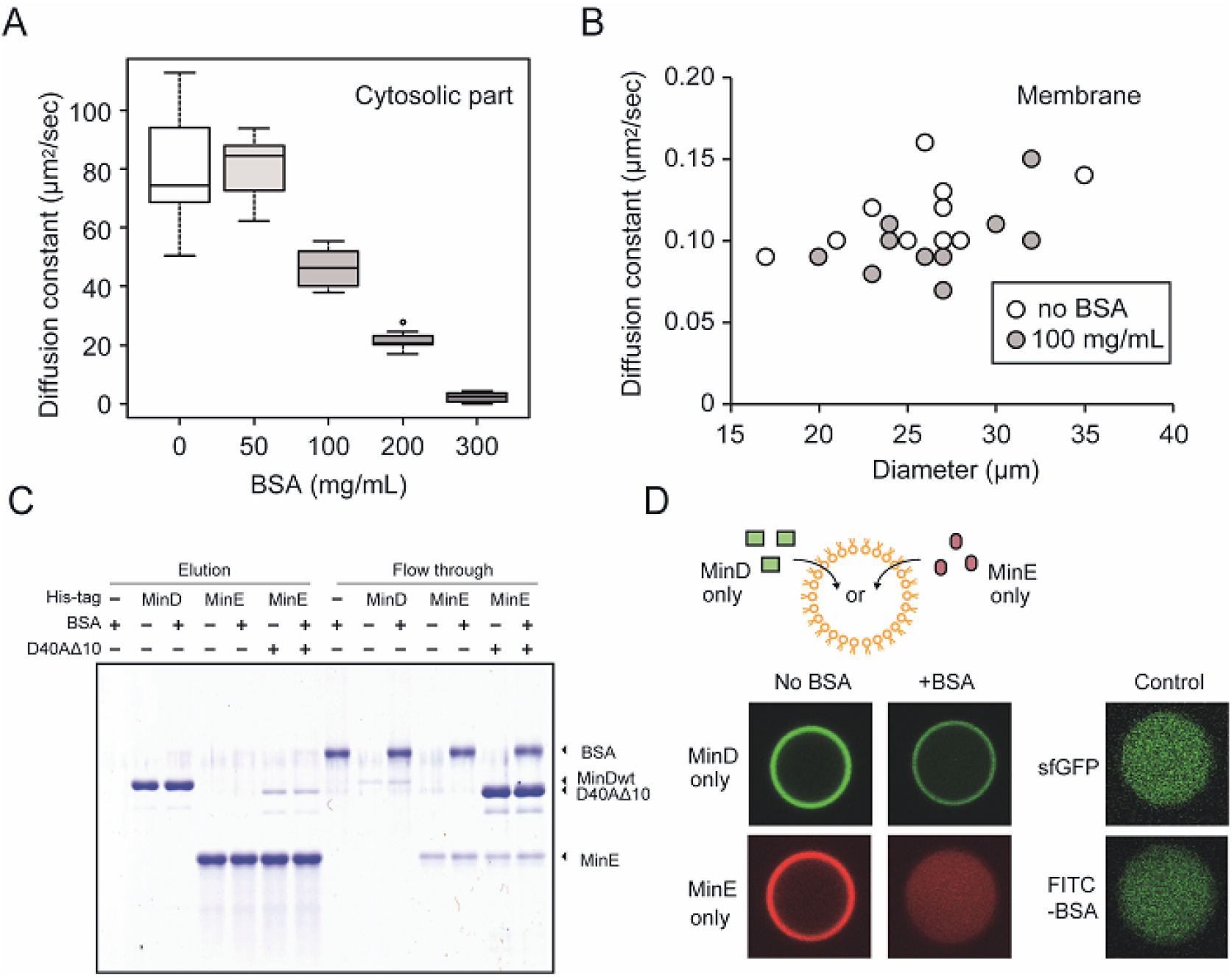
The effects of BSA on Min wave elements in cell-sized droplets. (**A**) Diffusion coefficients of sfGFP in solutions with various BSA concentrations entrapped in microdroplets (n = 10). (**B**) Diffusion coefficients of sfGFP-MinD attached on lipid membranes with or without 100 mg/mL BSA (n = 10) plotted as a function of microdroplet diameters. (**C**) Pull-down assay for BSA and Min proteins. His-sfGFP-MinD, MinE-mCherry-His, or sfGFP-MinD^D40A^Δ10 with MinE-mCherry-His was incubated with Ni-NTA resins under existence of BSA or not. The eluted fractions by imidazole and the flow through were visualized by CBB staining. (**D**) Inhibition of spontaneous binding between membranes and Min proteins by BSA. Either sfGFP-MinD or MinE-mCherry was encapsulated in the presence or absence of 50 mg/mL BSA. Microdroplets with 20 μm diameter were shown. As a control, the same experiments using 1 μM sfGFP only or 1 μM FITC-labelled BSA with 1 mg/mL BSA (right panel, 10 μm diameter droplets).

Next, to test the effect of BSA on reactions, we employed a pull-down assay to analyze the direct association of BSA with MinD or MinE. BSA was mixed with MinD or MinE immobilized on Ni-NTA beads through a histidine-tag. The pull-down assay showed that BSA flowed through the Ni-NTA with Min proteins, and therefore, no BSA band was found after eluting MinD or MinE by imidazole. These results indicated that BSA does not directly bind MinD or MinE. Similarly, the pull-down assay involving the use of MinD^D40A^Δ10 which kept binding with MinE due to its lack of ATPase activity (29), suggested that BSA does not enhance interaction of the MinDE complex (Figure 2C).

Finally, we found that BSA changes localization of MinE between cytosolic parts and on membranes. Each sfGFP-MinD and MinE-mCherry was encapsulated in micro-sized space fully covered with *E. coli* polar lipid, and localization of MinD and MinE was visualized using confocal fluorescence microscope. In the absence of BSA, almost all of the MinD and MinE were similarly localized on membranes. In contrast, BSA addition drastically changed the localization. In the presence of BSA, changes of MinD localization were relatively few, but localization of MinE on membranes completely disappeared (Figure 2D). Because sfGFP alone or BSA at low concentration (1 mg/mL) does not localize on membranes (Figure 2D), the spontaneous localization of MinE was not derived from a faulty membranes. These results suggested that changes in localization of MinE is a key feature for emergence of waves in microdroplets.

### Suppression of spontaneous membrane localization of MinE is the key to emergence of Min waves in micro-sized space

To quantify the details of MinE localization in microdroplets, we employed an index value for the localization ratio (c/m) obtained by dividing the concentration of MinE in cytosolic parts (c [1/μm^3^]) by those on membranes (m [1/μm^2^]) (Figure 3A). Both concentrations are expressed by characteristic concentrations in the cytosol, c_*b*_, and on the membrane, c_*s*_, such as c = *c*_0_*c*_*b*_ and m = *m*_0_*c*_*s*_, respectively. Here, we may freely choose the values of the characteristic concentrations, which would accordingly change the values of unitless concentrations *c*_0_ and *m*_0_. It is reasonable to assume that these quantities are proportional to the fluorescence intensity at the position at the cytosol *I*_*b*_ and on the membrane *I*_*s*_ with possibly different proportional constants as *c*_0_ = *α*_*b*_ *I*_*b*_ and *m*_0_ = *α*_*s*_*I*_*s*_, respectively. This argument ensures that the localization ration (c/m) is identified as c/m = (*α*_*b*_*c*_*b*_/(*α*_*s*_*c*_*s*_))*I*_*b*_/*I*_*s*_ with the ratio of fluorescence intensity up to a proportional constant. This argument implies that the relative value of c/m is a relevant quantity.

**Figure 3:**
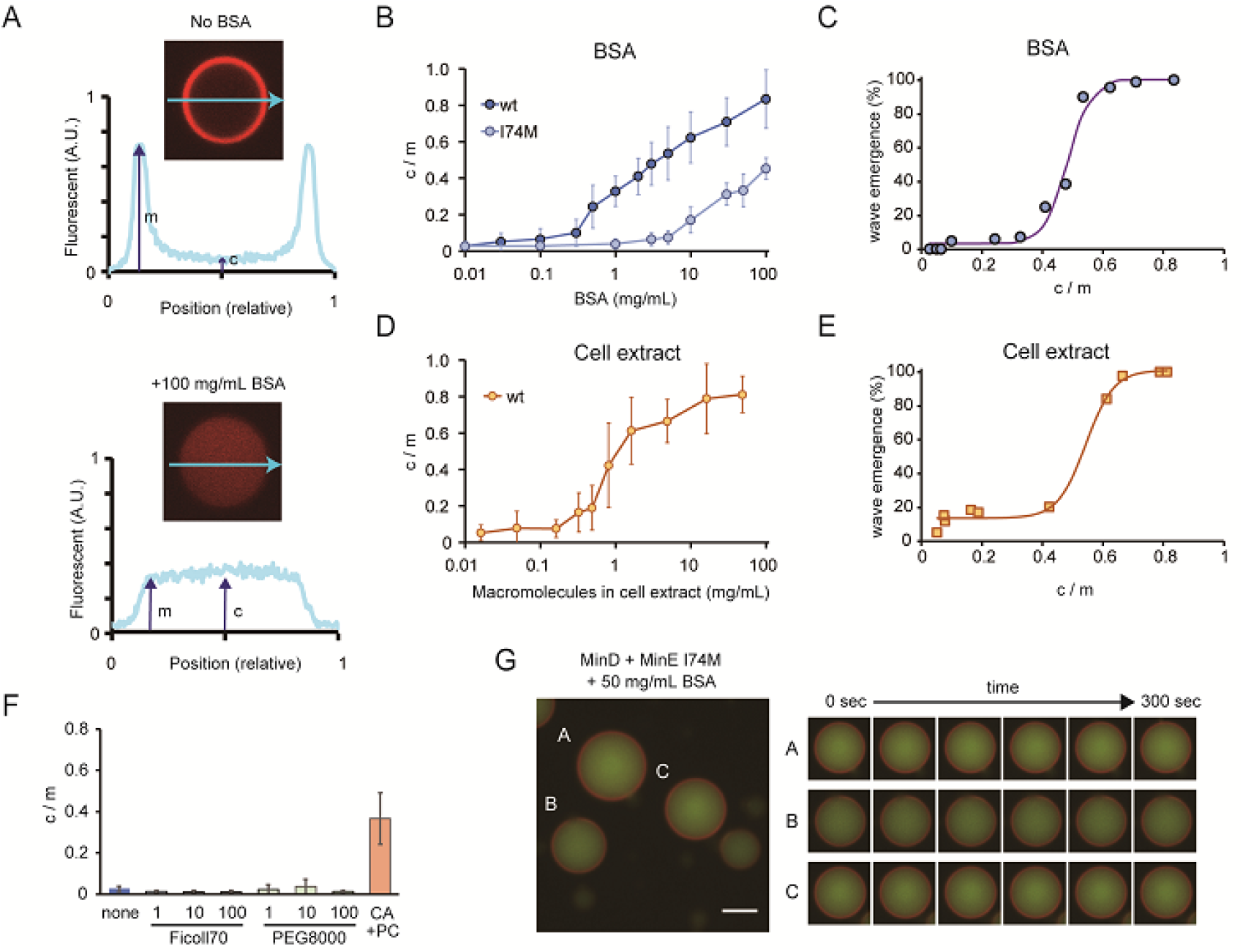
Relations between the rate of spontaneous localization of MinE on membrane and Min wave emergence. (**A**) Schematic illustrations of the method to evaluate MinE localization (c/m). (**B, D**) Changes of c/m of MinE-mCherry and its I74M mutants by various BSA concentrations (**B**) and by cell extract (**D**). (**C, E**) Probabilities of microdroplets with Min wave plotted as a function of c/m in the case of BSA (**B**) and cell extract (D), respectively. The fitting lines are sigmoidal curves. (**F**) Effects of macromolecular crowding agents (1, 10, 100 mg/mL) and the modified lipid condition (15% cardiolipin and 85% DOPC condition, abbreviated as CA+PC) on c/m of MinE-mCherry. Microdroplets smaller than 30 μm diameters were selected and c/m of 1 μM MinE-mCherry were evaluated. (**G**) Wide-view and sequential images of microdroplets encapsulating 1 μM sfGFP-MinD, 1 μM MinE^I74M^-mCherry, 2.5 mM ATP and 50 mg/mL BSA. Scale bar: 10 μm. In each figure, average and standard deviation were shown.

We measured fluorescence intensities at the center of microdroplets and the edges of signals. In this case, c/m becomes 1 when MinE is not localized on the membrane, whereas c/m becomes 0 when all MinE localize on membranes. As shown in Figure 3B, the c/m of MinE increased in proportion to BSA concentrations.

Then, we investigated the relation between c/m and the probability of Min wave emergence. Plots of wave emergence probability as a function of c/m controlled by BSA concentration showed that its relation is a sigmoidal as a threshold function (Figure 3C). Min waves were observed in a small fraction of microdroplets at c/m< 0.4 (<1 mg/mL BSA), and in almost all microdroplets at c/m>0.7 (>30 mg/mL BSA).

To check whether or not the effect is specific to BSA, we tested another protein crowder — a cell extract of *E. coli* prepared by sonication (24). In this case, we added an ATP recycling system to suppress ATP deletion due to the components of the cell extract. Addition of the cell extract modulate c/m of MinE was similar to the case for BSA, although its effect was stronger than BSA (Figure 4D). Moreover, the cell extract also led to the emergence of Min waves in the microdroplets covered with *E. coli* polar lipid extract (Video 7). The relation between c/m and the probability of Min wave emergence was similar to that of BSA (Figure 4E). These results indicated that high c/m is required for stable emergence of Min waves in a 3D closed geometry.

**Figure 4:**
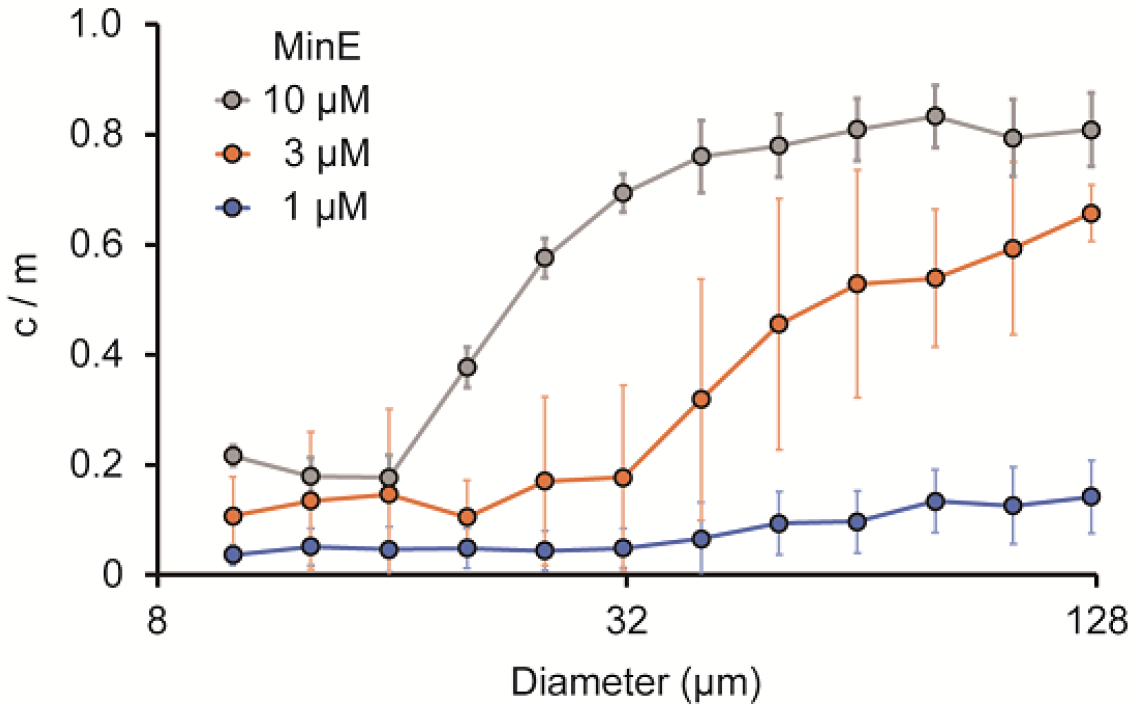
Size-dependence of c/m. Spontaneous localization of MinE-mCherry were plotted against sizes of microdroplets. Average of c/m ratio at each 0.1 logarithmic scale were shown (n = 208 for 1 μM, 386 for 3 μM, and 184 for 10 μM). Error bars indicate standard deviation.

Under conditions using macromolecular crowding reagents which do not lead to the emergence of Min waves (PEG8000 and Ficoll70), c/m was as low as similar to that without BSA (Figure 3F). Then, we checked c/m in the case of microdroplets covered with the modified lipid condition (15% cardiolipin and 85% DOPC), which causes Min waves without BSA. In the modified lipid case, c/m was near 0.4 (Figure 3F), which is as high as the minimal c/m value of BSA required for Min wave emergence. These results supported the notion that suppression of attachment of MinE on membrane without the aid of MinD is the key to emergence of Min waves in micro-sized space.

Experiments using a MinE mutant further supported the importance of the c/m. Recent studies have suggested that the conformation of MinE is in equilibrium between a free state of membrane targeting sequences (MTS) at the N-terminal (open conformation) and a packed structure (closed conformation). Open conformation preferably binds membranes, and several MinE mutants shift this equilibrium to the open state (14, 29). For an example, the I74M mutant of MinE stably maintains the open state, and the I74M mutant localizes on the membrane in *Δmin E. coli* cells, while wild-type MinE uniformly distributes in the cytoplasm (29). Our c/m analysis showed that the spontaneous membrane localization of I74M was less sensitive to BSA concentrations than that of wild type (Figure 3B). To match this result, no waves were observed even under 50mg/mL BSA conditions in the case of the I74M mutant (Figure 3G, Video 8).

### Space sizes of microdroplets changes the rate of spontaneous MinE membrane localization

As spatial factors to determine c/m of MinE, we can raise maximum levels of attachment on membranes and total amounts of MinE. In smaller microdroplets, the surface-area-to-volume ratio is large, and therefore, almost all MinE can localize on the membrane (meaning c/m ~ 0). By contrast, levels of MinE on membranes is near maximum levels for membrane localization in large spaces such as those found on 2D lipid bilayers, which derives a larger c/m. In fact, c/m in the case of 2D lipids, showing Min wave emergence in the absence of BSA, was estimated to be 0.76 (Figure 4—figure supplement 1). If this assumption true, c/m increases in proportion to sizes of microdroplets and its response to the space size is sensitive to MinE concentrations used.

To verify this point, we investigated the localization of MinE in various sizes of microdroplets in the absence of BSA. In small microdroplets with 10 μm diameter, c/m was near 0 and the value increased in higher concentrations of MinE. In larger microdroplets, c/m increased in proportion to the MinE amounts. Moreover, the increase of c/m was strongly dependent on MinE concentration. The diameter of microdroplets in which c/m reached 0.5 was approximately 45 μm at 10 μM MinE, and around 70 μm at 3 μM MinE (Figure 4). In the case of 1 μM MinE, c/m was maintained low within <130 μm (Figure 4). We should note that, in the presence of 10 mg/mL BSA, a minimum BSA concentration for Min wave emergence in microdroplets, c /m did not increase in size but was maintained high irrespective of droplet size (Figure 4—figure supplement 2). We also emphasize that at high concentration of MinE, Min wave did not emerge while MinE indicated high c/m ratio (Figure 1E, Figure 4). This suggests that high concentration of MinE inhibits wave emergence by other means, such as unbalanced MinDE ratios leading to a defect in typical turnover rates.

### Computational simulation for Min wave supports the importance of MinE localization for wave emergence

To understand the importance of MinE localization, we examined Min wave generation using computational simulations (see Methods and Appendix 1—3). We considered two models. Model I (Figure 5—figure supplement 1A) is simply based on the model proposed in (30) whereas Model II (Figure 5—figure supplement 1B) is based on a combination of the two models proposed in (30) and (31) to incorporate the effects of persistent MinE membrane binding and transformation from ADP-MinD to ATP-MinD in the cytosol (see Appendix 1). Based on these models, we investigated the effect of spontaneous MinE binding. This effect is characterized by the quantity, *c*_e,0_, which demonstrates the concentration of MinE on the membrane in the absence of MinD. To our knowledge, all the previous models lack this effect, that is, MinE was assumed to be in the cytosol without the presence of MinD (*c*_e_ = 0 when D_0_ = 0). This is because MinE in the absence of MinD fails to localize to the peripheral portion of the cell. This is in contrast with the observations of MinE binding on the membrane in the absence of MinD *in vitro* (29, 32, 33), and with our experiments demonstrating that MinE localization is a key factor to determine Min wave generation.

In both Model I and II, the concentration of MinE on the membrane becomes *c*_e,0_ in the absence of MinD. The rest of MinE is in the bulk of the cytosol, and therefore, c/m is given by (D_0_ − *αc*_e,0_) / *c*_e,0_ (see Appendix 2). Therefore, when *c*_e,0_ is smaller, c/m is larger.

First, we confirmed numerically that the rotating wave occurs in the closed membrane when MinE localization is weak, *c*_e,0_ ~ 0 (Video 9). The wave generation occurred when the total concentrations of MinD and MinE are comparable. We also observed that pole-to-pole oscillation occurs near the boundary between stationary state and rotating wave in the phase diagram of the two concentrations. The wave generation on the planar membrane was also observed for weak MinE localization, consistent with previous studies (30, 34).

Then, we studied the effect of spontaneous MinE localization on wave emergence using numerical simulations and linear stability analysis (see Methods). Figure 5 shows numerical results of the amplitude of the wave for the closed membrane in our models (red points for Model II and light red points for Model I). Irrespective to these models, the wave disappeared and the concentrations of MinD and MinE were uniform on the membrane when MinE localization is strong *c*_e,0_ ≫ 0. The condition of the wave generation may also be evaluated by linear stability analysis of the stationary state. In Figure 5, we showed by the (dark) shaded area the region at which the stationary state is linearly unstable. In this area, Min waves occurred. Both the numerical results and linear stability analysis provided evidence that above *c*_e,0_ = 0.03 the Min wave disappears.

**Figure 5:**
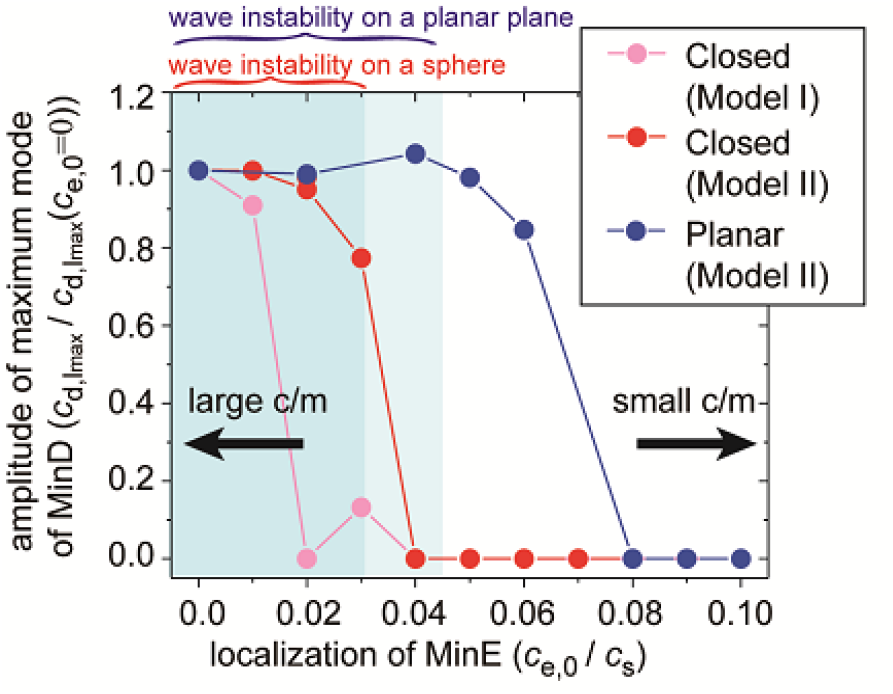
Simulation results of wave generation in the presence of membrane attachment of MinE. The simulation results of wave generation in the presence of spontaneous MinE interaction at 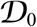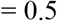 and 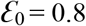 (see Methods) are shown. The results of the closed spherical membrane and the planar membrane are shown. Waves are characterized by the amplitude of the first mode (*l* = 1) in spherical harmonics expansion of the closed membrane, and by the maximum amplitude at a finite wave number in the planar membrane. The amplitude is normalized by its value without spontaneous MinE localization. Stability analysis of the homogeneous state calculated by the real part of maximum eigenvalue shows that the homogeneous state is stable in the dark (closed membrane) and light (planar membrane) shaded areas, but linearly unstable otherwise.

Consistent with our experimental results, numerical simulations indicated that the degree of spontaneous MinE binding shifts the conditions for Min wave emergence (Figure 5). We also performed the same analysis for the planar membrane. The linear stability analysis (light shaded area) in Figure 5 showed that the critical concentration of MinE localization is higher in the planar membrane. Furthermore, the numerical results illustrated an even larger shift of the transition point, as shown in blue points in Figure 5. These results suggested that the condition is dependent on the size of the membrane; under confinement, the shift is sufficiently strong to eliminate wave generation at stronger MinE localization. On the other hand, wave generation of the planar membrane was less suppressed, and thus, it remained at stronger MinE localization on the membrane.

### Theoretical analysis reveals that confinement regulates Min wave emergence

To investigate the effect of confinement, we studied the two models introduced above (Model I and II). We used these two models because they incorporate the two effects (persistent MinE membrane binding and transformation from ADP-MinD to ATP-MinD in cytosol), which were assumed to play essential roles in the wave generation, but have been studied separately. We found that in the closed geometry, these two models reproduce the same results, suggesting that under confinement the difference between the models is not important.

Figure 6A shows the phase diagram of the wave generation in the total MinD and MinE concentrations according to the linear stability analysis of Model II. The MinE wave occurred when both the concentrations were above the values at the phase boundary for the first mode (*l* = 1). The condition of the stability of the stationary state is dependent on the spatially inhomogeneous modes. The mode number is denoted by *l*. The zeroth mode (*l* = 0) expressed uniform concentration on the membrane whereas the first mode (*l* = 1) expresses inhomogeneous distribution with one wavelength on the membrane (see the inset in Figure 6A). The homogeneous oscillation (*l* = 0), in which the concentrations of MinD and MinE are uniform on the membrane but oscillate in time, occurs at another phase boundary shown in Figure 6A. The phase boundary of the homogeneous oscillation requires higher concentrations than that of the Min wave, resulting in wave generation rather than uniform oscillation in the closed membrane. This behavior is not obvious in reaction diffusion systems. For any two-variable reaction-diffusion equations, it can be shown that uniform oscillation occurs rather than the wave of the first mode (35). In our models, wave generation did occur by additional degrees of freedom.

**Figure 6:**
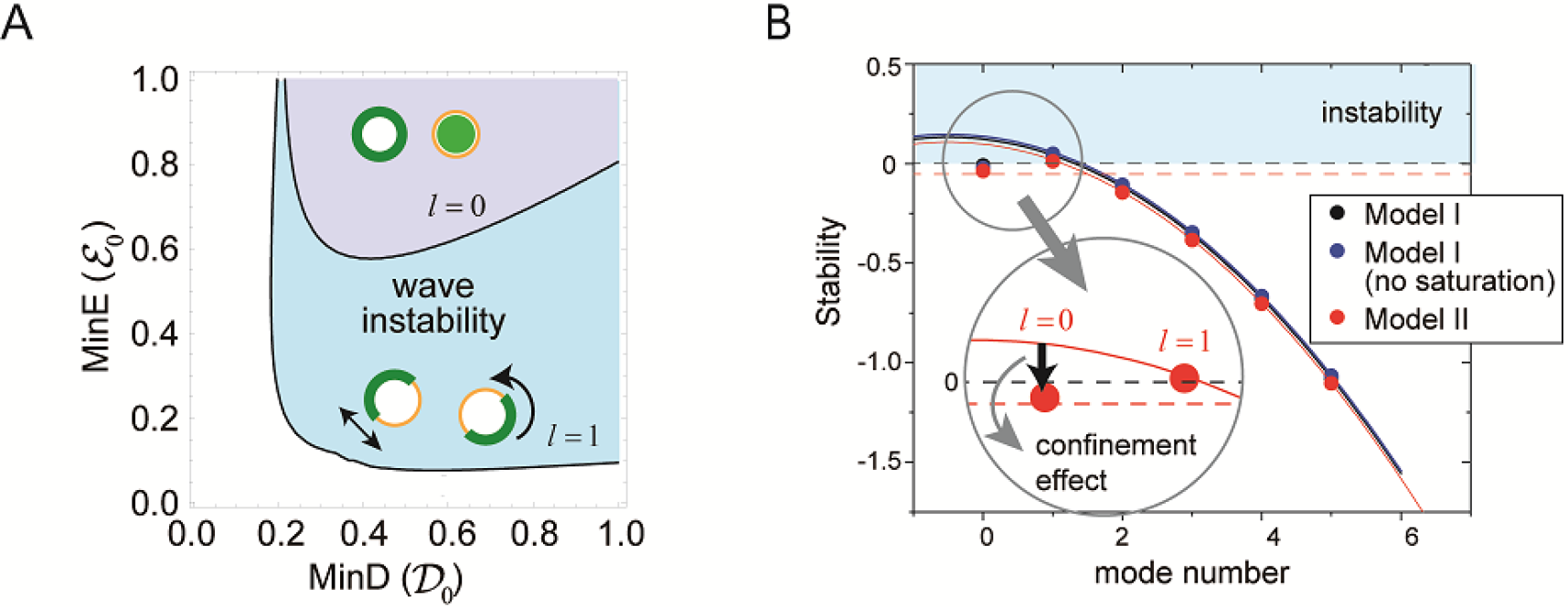
Regulation of generation and stability of Min wave by confinement. (**A**) The phase diagram obtained from linear stability analysis of wave instability (the first mode, *l* = 1) and homogeneous oscillation (the zeroth mode, *l* = 0) under various total concentrations of MinD and MinE in the closed membrane of its size *R* = 5 in Model II. (**B**) Stability of the homogeneous state for each mode obtained by a real part of eigenvalue in different models near the transition point of wave generation, 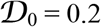 and 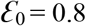. Instability is demonstrated by positive eigenvalues. The black dashed line indicates neutral stability in which the real part of eigenvalue is zero. The theoretical results under approximation that neglect the effect of bulk dynamics are demonstrated by the solid lines (See Methods and Figure 6—figure supplement 1). Each color (black, blue, and red) corresponds to a different model. The dashed red line shows the stability of the homogeneous state theoretically obtained by including the effect of confinement.

To investigate theoretically the mechanism of suppression of the uniform oscillation resulting in inhomogeneous wave generation, we considered the generic frame work to combine the two models outlined in Appendix 4. Our method enabled us to eliminate the bulk cytosol concentration field. The condition of the wave generation was identified by the real part of the largest eigenvalue Re *σ* > 0, where the eigenvalues, *σ*, were then obtained by solving the following equation:

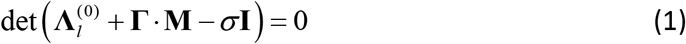

All the terms in the determinant are *n*×*n* matrices under *n* concentration fields on the membrane. The first term describes the reaction on the membrane whereas the second term expressed the effect of bulk cytosol. Here, **I** denotes the *n*×*n* identity matrix and **Γ** shows the coupling of the reactions on the membrane with the bulk cytosol concentrations close to the membrane (see Appendix 4). The effect of confinement in **M** appears from its dependence on the size of the system, such as the radius *R* of sphere or the height *H* of the bulk on the planar membrane. Figure 6B shows the real part of the largest eigenvalue for the closed membrane as a function of the number of modes. The eigenvalue is positive only at the first mode, suggesting that wave instability occurs instead of uniform oscillation. This result is independent of choice of Model I or II, and furthermore independent of the saturation term (see Appendix 3).

From the theoretical analysis, we were able to identify three effects of confinement: (i) The homogeneous stationary solution is dependent on the system size through *α*, (ii) the diffusion on the membrane inhibits higher-mode (smaller length scale) inhomogeneity (see Eq.(32)), and (iii) the effect of the dynamics of the bulk concentrations in Eq.(3) modifies the stability. Among the three contributions, the second one is easily computed once we know the eigenvalues at the zero mode for the matrix

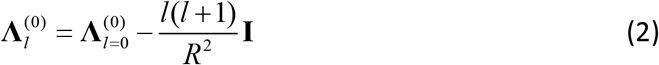

In Figure 6B, this is demonstrated by the solid lines for each model (see also Figure 6—figure supplement 1). It is evident that the stability at the higher modes is dominated by this effect. On the other hand, the eigenvalue at the zero mode is deviated from the lines. This result is explained by the effect of (i) and (iii), suggesting that the mechanism of the wave instability is oscillatory instability at the first mode (*l* = 1) with suppression of instability at the zero mode (*l* = 0) due to the effect of confinement.

To see more details about the effect of confinement, we investigated the second term in Eq.(1)(Figure 7). For the spherical membrane (Figure 7A), the effect of confinement is given by

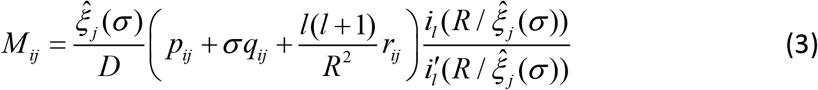

where

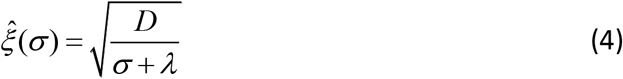

and *i*_*l*_(*x*) is *l* th-order of the modified spherical Bessel function of the first kind and *i* ′_*l*_(*x*) = *di*_*l*_(*x*) / *dx*. This effect is significantly different from the planar membrane (Figure 7B), in which the effect of bulk is expressed by

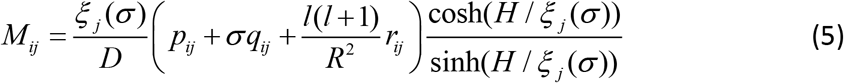

where the length scale *ξ* is expressed by the eigenvalue, *σ*, and wave number, *k* = |**k**|, in (10) as

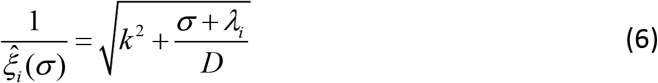

**Figure 7:**
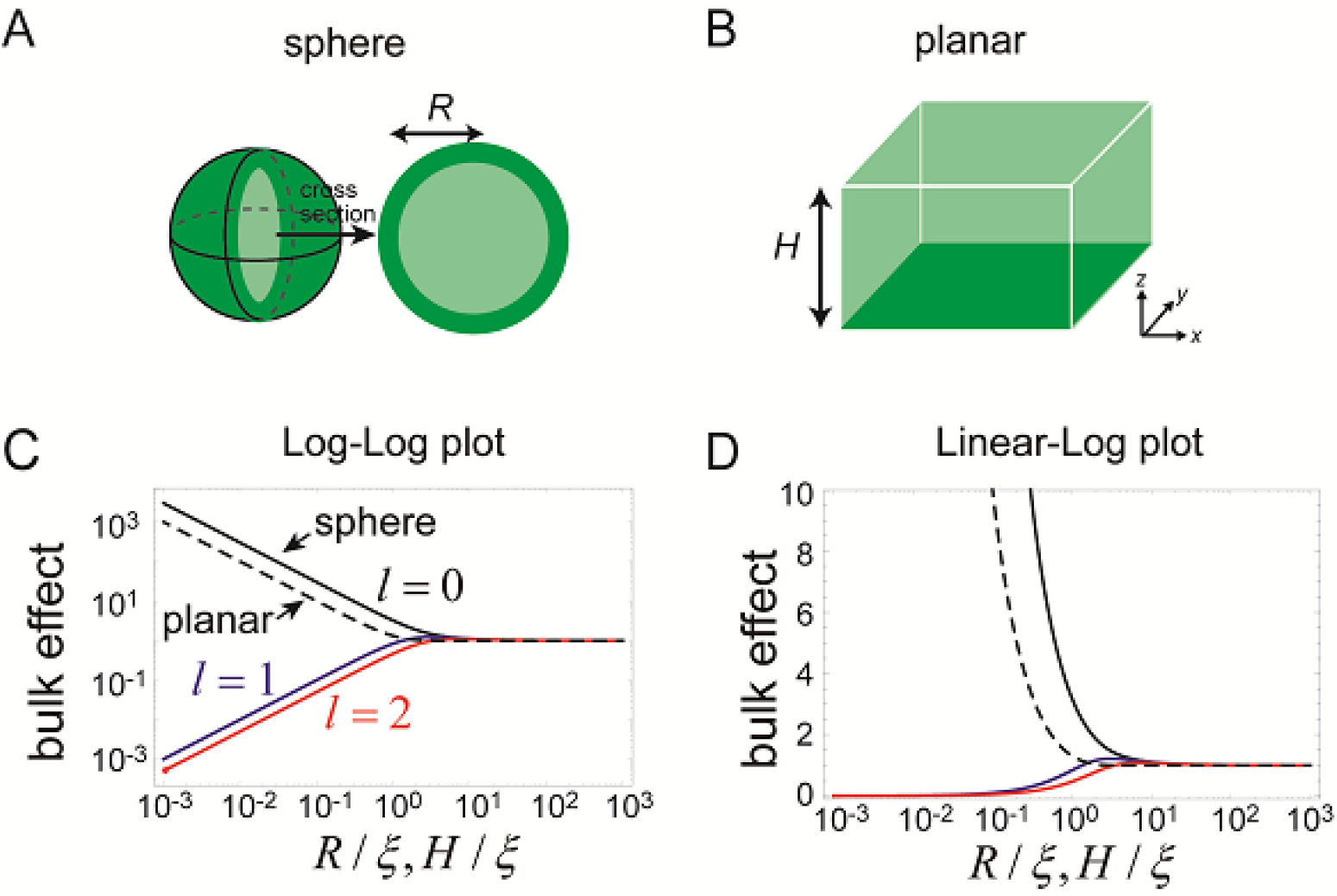
Effects of bulk in the models. (**A, B**) Schematic illustration of closed membrane with the radius, *R*, (**A**), and planar membrane with the height, *H*, (**B**). Membrane-bound proteins are denoted by dark green, while bulk proteins are shown in light green. (**C, D**) The log-log (**C**) and log-linear (**D**) plots of 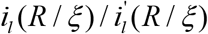 as a function of *R* / *ξ*. The homogeneous mode (*l* = 0, black) and the two lowest inhomogeneous modes (*l* = 1, blue, and *l* = 2, red) are shown in solid lines. The corresponding term in the planar membrane, cosh(*H* / *ξ*) / sinh(*H* / *ξ*) as a function of *H* / *ξ*, is shown in dashed line.

The magnitude of this effect is represented by 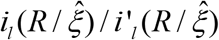 for the closed membrane and 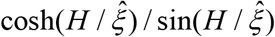 for the planar membrane, both of which are shown in Figure 7C and D. As the size *R* and *H* decreases, the effect becomes stronger for the zero mode of the spherical membrane and for all the wave numbers of the planar membrane. This result is in contrast with the higher modes (*l* ≥ 1) of the spherical membrane. Thus, for a small system, the effect of the dynamics of bulk remains only for the zeroth mode of the spherical membrane. The physical picture of this result is that, an inhomogeneous concentration associated with the higher-order modes is suppressed in a small system, while in the planar membrane, inhomogeneity in the plane may exist independently from the direction perpendicular to the membrane.

### Excitability may occur in the planar membrane

Our numerical simulations suggested that the robustness against MinE localization is stronger than the prediction by the linear stability analysis (Figure 5). One possible reason is that the wave generation is dependent on initial perturbation of the concentration fields due to the excitability of the system. The homogeneous stationary state is linearly stable but responds largely against finite perturbation (Figure 8). In fact, the instability of the planar membrane starts from a core of wave emergence rather than uniform oscillation on the planar membrane, as observed in a previous study(7).

**Figure 8:**
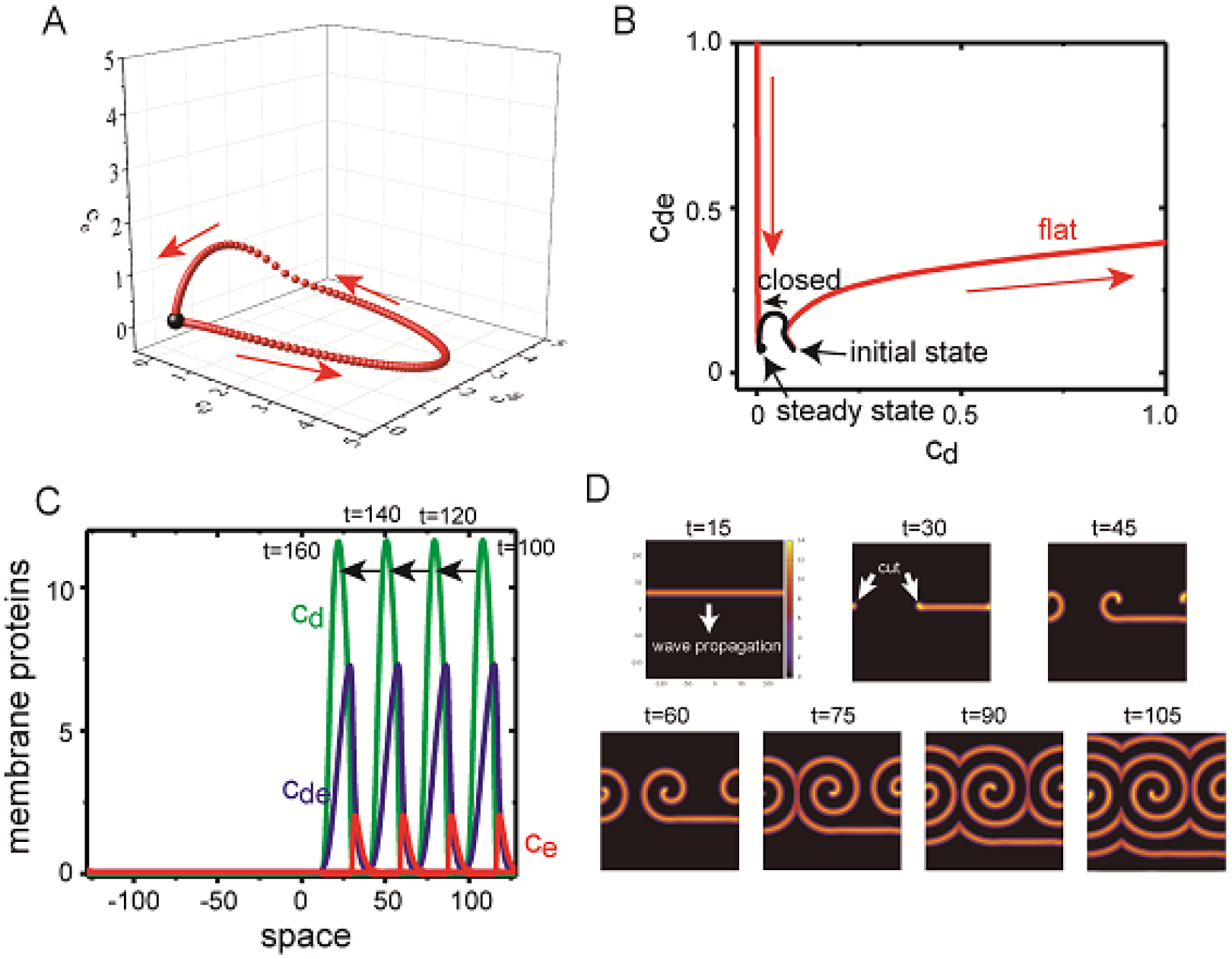
Excitability of Min waves on the planar membranes. (**A**) Trajectory of the concentrations of membrane proteins in (*c*_*d*_, *c*_*de*_, *c*_*e*_) coordinates starting from the initial condition slightly shifted from the homogeneous stationary state at 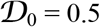, 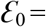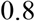 and *c*_*e*,0_ = 0.07. (**B**) Trajectory near the stationary state in (*c*_*d*_, *c*_*de*_) plane on the planar membrane (*H* = 256, red) and on the closed membrane (*R* = 5, black). (**C**) A propagating pulse in one-dimensional membrane surrounded by two-dimensional bulk. (**D**) A propagating band and a spiral wave in two-dimensional planar membrane underneath the three-dimensional bulk. Initially, an isolated band is prepared and let it propagates, and then cut it at *t* = 22.5 to make a spiral wave.

To investigate excitability of the planar membrane under Model II, we first studied the dynamics of the concentrations of membrane-bound proteins without diffusion on the membrane; namely, the bulk concentration was one dimension, and the membrane concentration was zero dimension. At *c*_*e*,0_ = 0.07 in which the homogeneous stationary state was linearly stable, an initial condition of *c*_*d*_ was shifted from the value at the stationary state 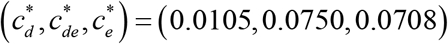 while 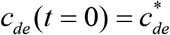 and 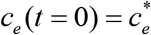. When the deviation, 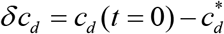, was small, the system quickly returns to the stationary state. When *δc*_*d*_ > 0.075, the system initially went away from the stationary state and exhibited a completely different trajectory (Figure 8A and B). This behavior suggests that the system is excitable, in which the system is stable against a small perturbation but responds largely against a perturbation above a particular threshold. In contrast with the planar membrane, the closed membrane did not show excitability (Figure 8B), and the system quickly relaxed to its stationary state without travelling in a large path. It is known that excitable systems may exhibit dissipative solitary pulses propagating in one direction with fixed speed, and spiral and turbulent waves in two dimensions (36, 37). In fact, Model II demonstrated a propagating solitary wave when the initial condition was chosen appropriately in one- (Figure 8C) and two-dimensional (Figure 8D) membranes. A spiral wave was obtained by cutting a solitary band in the two-dimensional membrane (Figure 8D), a phenomenon which has also been observed in other excitable systems (38).

### Early stage of Min wave emergence in microdroplets

Finally, we analyzed the early stage of Min wave emergence in micro-sized space (Figure 9, Video 10). In the case of small microdroplets that only show a single wave inside, time-lapse imaging of Min proteins showed that pulsing between cytosolic parts and membrane surface is the initial stage of Min wave emergence, similar to a previous report(12). However, our imaging demonstrated that the pulsing pattern transforms to pole-to-pole oscillation, and then, settles in traveling waves. This transition of wave patterns is not specific to wet experiments but can be recapitulated by our computational simulation (Figure 9, Video 9). Introduction or reduction of stochastic noises to the simulation did not change the results, indicating that this transition proceeds in a deterministic manner; namely, wave instability underlying reaction-diffusion coupling is the only driving force to lead to the emergence of Min waves in cell-sized space, showing the specificity of the wave emergence mechanism in such space.

**Figure 9:**
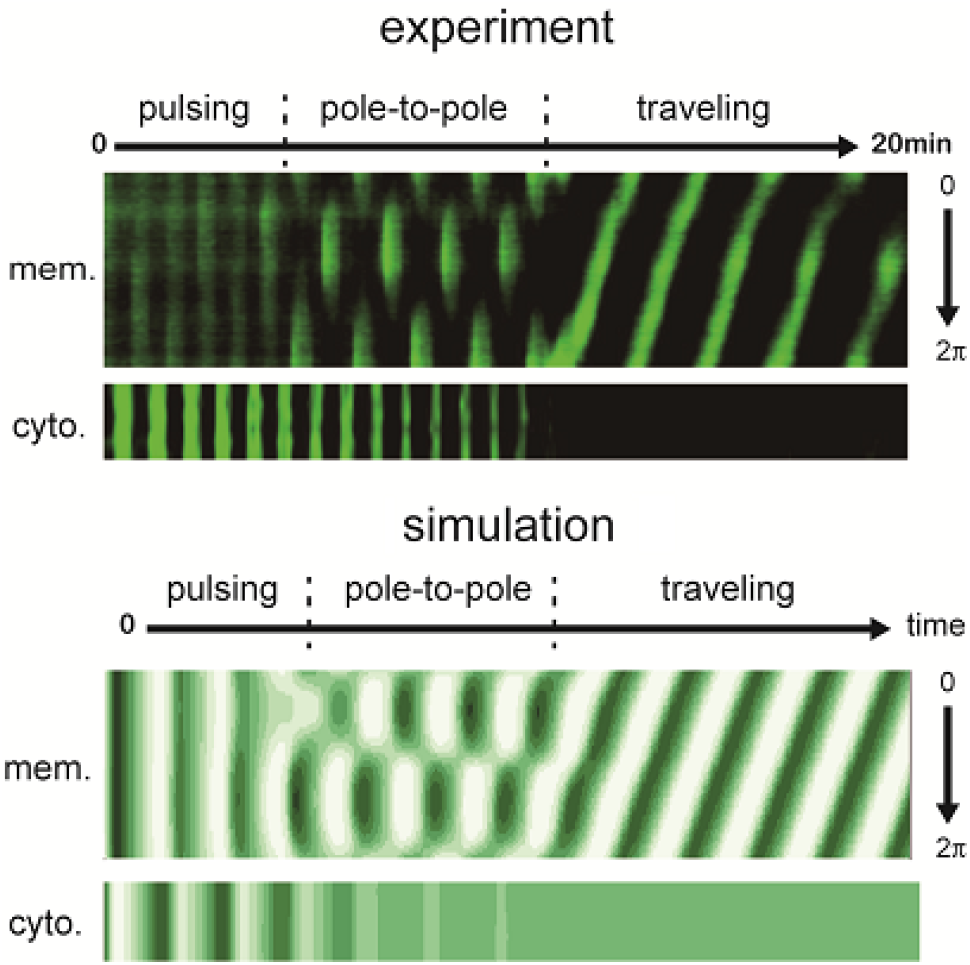
Early stages of Min wave emergence inside microdroplets and on 2D membranes. Transition of patterns from pulsing during the initial stage of wave emergence to stable traveling wave. Kymographs of sfGFP-MinD in the membranes and inner media of droplets obtained by experiments (**top**) and numerical simulation without noise (**bottom**) are shown.

## Discussion

Conditions for Min wave appearance have been regarded as the same between open systems such as on 2D planar membranes and closed geometry as in fully confined cell-sized space. In this study, we show that the conditions for Min wave appearance are limited in cell-sized closed space as compared with the case of on a 2D planar membrane. From experiments and simulation, it has been shown that the rate of spontaneous localization of MinE is an important factor to determine generation of Min waves in cell-sized closed space.

Due to the large surface-area-to-volume in the cell-sized space, even for weak interactions, localization to the membrane becomes prominent compared to the flat membrane system. Spontaneous localization of MinE on membranes works in an inhibitory manner with respect to the generation of Min waves, but it is suppressed in the presence of a protein crowder such as BSA or cell extract to generate Min waves. This effect is observed at a relatively low concentration (1–10 mg/mL) than the case of crowding cytoplasm in living cells (100–300 mg/mL) and is not observed with synthetic polymers such as PEG8000 and Ficoll70. Hence, BSA and cell extract are considered to modify the interaction between MinE and membranes, and the mechanism is different from the effect known as crowding, such as increasing viscosity.

Because a previous report has indicated that BSA at high concentration (>10mg/mL) attaches to the lipid membrane(39), and several proteins in cell extract are assumed to interact with such membranes, a plausible explanation of the effect of the protein crowders is competitive inhibition. To match this assumption, tuning lipids conditions to reduce spontaneous MinE attachment on membranes (Figure 3F) seems to be important for generation of Min waves without aid by auxiliary molecules, as reported previously (12, 15). Although estimation of the exact strength of interaction between MinE and membranes in cell-sized space is important for understanding the details of spontaneous membrane binding of MinE, we failed to do that due to the technical difficulties associated with the measurement. However, the level is assumed to be weak from a previous study using 2D planar membranes (1/100 of the strength of MinD binding) (33).

Another possibility to suppress the membrane attachment of MinE on membranes is regulation of the open-closed equilibrium state by excluded volume or other effects. This point will be clarified by analyzing the open-closed equilibrium state of MinE in a similar manner to that for a previous study (29) in the presence of protein crowders.

We should note that the levels of spontaneous localization of MinE is not the only determinant factor of Min wave generation. In the I74M mutant, high concentrations of BSA suppress the spontaneous localization of MinE, but no Min waves were observed. Because I74M mutation is assumed to fix the open state of MinE, other parameters in reaction kinetics should change. As a result, the generation of Min waves is suppressed. In fact, I74M mutants do not reproduce Min waves even in living cells (29). The effects of MinE conformation shifts should be clarified in the future by analysis using various MinE mutants. However, it is plausible that spontaneous localization of MinE to the membrane is a direct regulator of Min waves in cell size spaces, as demonstrated in this study.

Our computational simulations showed that the condition for Min wave emergence depends on membrane MinE accumulation and membrane size (Figure 5). Cell-sized space stabilized homogenous state (Figure 6), and therefore, Min waves, oscillation inhomogeneous in space, emerge instead of homogeneous oscillation. In contrast, for the flat membrane, the wave robustly appears against the increase of the spontaneous MinE binding. Our simulation suggested that this robustness is originated from the coupling between membrane and bulk dynamics, and the excitability of the system. If the system is excitable, the homogeneous stationary state is linearly stable, but responds largely against finite perturbation (Figure 8). Although our simulation suggests that the effects by excitability is stronger than space size effects, it is still elusive whether or not the excitability shown by this Min wave is model independent. This question would be clarified by investigation of the wave generation under controlled initial conditions in further experiments.

Recent *in vitro* reconstitution studies have demonstrated that biosystems in cell-sized space show characteristic features of those biosystems not found in test tubes. For example, cell-sized space enhances formation of the actomyosin ring (40), affects aqueous phase separation (16, 41), and confers scaling properties of spindle shapes (42). Although these studies have determined that space size is a cue to change the behaviors of biosystems, biochemical parameters and mechanisms underlying their behavior have been assumed to be equal irrespective of space sizes. Our present study provides evidence that cell-sized space shifts the equilibrium of membrane binding of proteins, and changes conditions for generation and stability of iRD waves from those in 2D planer membranes. As theoretical analysis of Min wave behaviors have indicated, the control of generation and stability by cell-sized confinement are expected to be universal features among iRD waves.

Furthermore, equilibrium shifts of protein localization by surface-to-volume effects should be universal among biosystems because maximum attachment levels and total amounts of the factor explained the shift. These points will be elucidated by *in vitro* reconstitution of other iRD systems (1–4).

## Materials and Methods

### Expression and purification of MinD and its mutant

In this study, all *Escherichia coli* cells were cultivated in LB medium. To construct pET15b-MinD, MinD gene was cloned from *E. coli* MG1655 genome by PCR into pET15b (Merck Millipore, Billerica, MA, USA) by Gibson assembly (New England Biolabs, Ipswich, MA, USA). To construct pET15-sfGFP-minD, sfGFP gene amplified from pET29-sfGFP (43) by PCR were cloned into N-terminal of MinD by Gibson assembly. To construct pET15-sfGFP-MinD^D40A^Δ10, D40A mutation and deletion of C-terminal 10 amino acids were introduced into pET15-sfGFP-minD by using the PrimeSTAR Max mutagenesis protocol (TaKaRa, Shiga, Japan). Similarly, K11A mutation was introduced into pET15-sfGFP-minD to construct pET15-sfGFP-MinD^K11A^. *E. coli* BL21-CodonPlus(DE3)-RIPL (Agilent Technologies, Santa Clara, CA, USA) cells were transformed with the resultant plasmids.

Proteins were expressed by 1 mM IPTG at OD_600_=0.1-0.2 and further cultivation at 37°C for 3 to 4 h. The cells were collected by centrifugation and suspended in LS buffer [50 mM NaH_2_PO_4_ (pH 7.6), 300 mM NaCl, 10 mM imidazole, 10 mM dithiothreitol (DTT), 0.1 mM phenylmethylsulfonyl fluoride (PMSF)] with 0.2 mM ADP-Mg. The collected cells were disrupted by sonication using a Sonifier250 (Branson, Danbury, CT, USA), and the supernatant of the crude extract was fractionated by centrifugation at 20,000g at 4°C for 30 min. To purify His-tagged proteins, the crude extracts mixed with cOmplete His-Tag purification resin (Roche, Basel, Switzerland) were loaded onto a polyprep chromatography column (Bio-Rad, Hercules, CA, USA), and washed with 25 mL WS buffer [50 mM NaH_2_PO_4_ (pH 7.6), 300 mM NaCl, 20 mM imidazole, 10% glycerol, 0.1 mM EDTA, and 0.1 mM PMSF]. His-tagged proteins were eluted with EL buffer [50 mM NaH_2_PO_4_ (pH 7.6), 300 mM NaCl, 250 mM imidazole, 10% glycerol, 0.1 mM EDTA, and 0.1 mM PMSF]. EL buffer was exchanged with storage buffer [50 mM HEPES-KOH (pH 7.6), 150 mM GluK, 10% glycerol, 0.1 mM EDTA] with 0.2 mM ADP-Mg by ultrafiltration using AmiconUltra-15 10k and AmiconUltra-0.5 30k filters (Merck Millipore).

For pull-down assay, His-sfGFP-MinD^D40A^Δ10 was treated with thrombin (Wako, Osaka Japan) in the storage buffer at 4°C overnight. Then, the cleaved His-Tag (2kDa) was removed from the sfGFP-MinD^D40A^Δ10 (55kDa) solution by ultrafiltration using AmiconUltra-0.5 50k filters (Merck Millipore). Proteins in the storage buffer were stored at −80°C. Protein purity and concentrations were estimated by Comassie Brilliant Blue (CBB) staining after separating by sodium dodecyl sulphate polyacrylamide gel electrophoresis (SDS-PAGE) and bicinchoninic acid (BCA) assay.

### Expression and purification of MinE and its mutant

To construct pET29-minE-His and pET29-minE-mCherry-His, MinE and mCherry genes were amplified from the *E. coli* K12 MG1655 genome or the pET21b-RL027A (Addgene, Cambridge, MA, USA), respectively, and were cloned into pET29a (Merck Millipore) by Gibson assembly. 6xHis-tag at C-terminal of MinE or mCherry was attached by PCR. The I74M mutation of MinE was introduced by using pET29-MinE-mCherry and PrimeSTAR Max mutagenesis protocol. *E. coli* BL21-CodonPlus(DE3) RIPL cells were transformed with the resultant plasmids.

Proteins were expressed by 1 mM IPTG at OD_600_=0.1-0.2 and further cultivation at 37°C for 3 to 4 h (pET29-minE-His) or at 16°C for 12h (pET29-minE-mCherry-His and its mutant). Cells were collected by centrifugation, resuspended in LS buffer, and purified using the same protocol as described for MinD. The elution fraction of MinE-mCherry-His diluted 5-to 10-fold with HG buffer [50 mM HEPES-KOH, pH 7.6, 10% glycerol, and 0.1 mM EDTA] was further purified by using Hitrap Q HP column (GE Healthcare, Chicago, IL, USA) and AKTA start (GE Healthcare). Briefly, the diluted fraction was loaded onto the column equilibrated with A buffer [50 mM HEPES-KOH (pH 7.6), 50 mM NaCl, 10% glycerol, and 0.1 mM EDTA], and washed using the same buffer. Proteins were eluted by IEX protocol of AKTA start using A buffer and B buffer [50 mM HEPES-KOH (pH 7.6), 1 M NaCl, 10% glycerol, and 0.1 mM EDTA]. Peak fractions monitored by SDS-PAGE were collected and exchanged with the storage buffer using AmiconUltra-15 10k and AmiconUltra-0.5 10k filters (Merck Millipore). Samples were stored at −80°C, and protein purity and concentrations were estimated by CBB staining after separating by SDS-PAGE and BCA assay. For MinE-mCherry-His proteins, concentrations were estimated by quantitative CBB staining using Fiji software (National Institutes of Health, Bethesda, MD, USA) to avoid signal contamination from mCherry absorbance.

### Expression and purification of MinC-sfGFP

MinC gene and sfGFP gene were amplified and cloned into the pET15b vector by the same procedure for MinD. *E. coli* BL21-CodonPlus(DE3)-RIPL cells were transformed with the resultant plasmid. IPTG was added at OD_600_=0.1-0.2 to 1 mM, and cells were further cultivated at 16°C overnight. The protocol for purification, storage, quantification of MinC-sfGFP was the same as for MinD except no ADP-Mg addition.

### Preparation of E. coli cell extract

*E. coli* BL21-CodonPlus(DE3)-RIPL cells were cultured in LB medium at 37°C. Cells at OD_600_ = 0.7 were collected by centrifugation and suspended in LSE buffer [25 mM Tris-HCl (pH 7.6), 250 mM NaCl, and 10 mM GluMg]. Then, cells were disrupted by sonication using a Sonifier250, and the supernatant of the crude extract after centrifugation at 30,000g for 30 min at 4°C were collected as cell extract. To remove genome DNA and RNA, cell extract was incubated at 37°C for 30 min. The supernatant after centrifugation at 30,000g for 30 min at 4°C was exchanged with the RE buffer [25 mM Tris-HCl (pH 7.6), 150 mM GluK and 5 mM GluMg] using AmiconUltra-15 3k and AmiconUltra-0.5 3k filters (Merck Millipore). The sample was stored at −80°C, and protein concentration was estimated by BCA assay. Concentrations of RNA such as ribosomal RNA, tRNA, and mRNA were estimated by 260 nm absorbance. Macromolecule concentrations were determined by the summation of protein and RNA concentration (44).

### Preparation of supported lipid bilayers (SLBs) on a mica layer

The general protocol was followed according to a previous report (11). *E. coli* polar lipid extract (Avanti, Alabaster, AL, USA) in chloroform at 25 mg/mL was dried by argon gas flow. The lipid film was further dried in a desiccator for at least 30 min at room temperature, followed by resuspension in TKG150 buffer [25 mM Tris-HCl (pH 7.6) and 150 mM GluK] to a lipid concentration of 5 mg/mL and then gentle hydration at 23°C for at least 1 h. The lipid solution was then vortexed for 1 min and sonicated using a Sonifier250 for 10 min to 15 min (Duty10%, Output1) to obtain small unilamellar vesicles (SUVs). SUV solution was diluted to 2 mg/mL with TKG150 buffer, and CaCl_2_ was added to a final concentration of 0.1 mM. This solution was applied to a thin mica layer mounted on the bottom of a glass base dish (Iwaki, Tokyo, Japan). After a 1-h incubation at 37°C, excess SUVs were washed with RE buffer.

### Self-organization assay for Min proteins on SLBs

For the self-organization assay, a reaction mixture containing 2.5 mM ATP, 1 μM His-sfGFP-MinD, and 1 μM MinE-mCherry-His in RE buffer was added to the SLBs, followed by incubation at room temperature for 10 min prior to microscopic observation. Self-organization of Min proteins was observed using a fluorescent microscope (Axiovert 200M; Carl Zeiss, Jena, Germany) with a CMOS camera using an ORCA-Flash4.0 V2 (Hamamatsu Photonics, Shizuoka, Japan) or a confocal laser-scanning microscope FV1000 (Olympus, Tokyo, Japan).

### Self-organization assay inside lipid droplets

The general protocol for microdroplets preparation was followed according to a previous report (22). *E. coli* polar lipid extract (Avanti) in chloroform at 25 mg/mL was dried by argon gas flow and dissolved with mineral oil (Nacalai Tesque, Kyoto, Japan) to 1 mg/mL in glass tubes. The lipid mixture was then sonicated for 90 min at 60°C using Bransonic (Branson). For preparation of the modified lipid mixture, 15% of 10 mg/mL *E. coil* Cardiolipin (CA) (Avanti) and 85% of 10 mg/mL 1,2-dioleoyl-sn-glycero-3-phosphocholine (DOPC) (Avanti) dissolved in chloroform were mixed and microdroplets were prepared as the same way for *E. coli* polar lipid extract. For the self-organization assay, the reaction mixture consisted of 1.0 μM His-sfGFP-MinD, 1.0 μM MinE-mCherry-His, 2.5 mM ATP, and macromolecules [BSA of Cohn Fraction V (A6003, Sigma-Aldrich, St. Louis, MO, USA), *E. coli* cell extract, Ficoll70 (Santa Cruz Biotechnology, Dallas, TX, USA), or PEG8000 (Promega, Madison, WI, USA) in RE buffer]. Concentrations of BSA and *E. coli* cell extract were varied to evaluate the concentration dependence of Min waves. To avoid depletion of ATP due to endogenous enzymes in *E. coli* cell extract, 80 mM creatine phosphate and 0.4 mg/mL creatine kinase were added for the assay using cell extract. The reaction mixture (2 μL) was added to the lipid mixture (100 μL), and lipids microdroplets were obtained by emulsification with tapping. A portion of the mixture (15μL) was gently placed into two glass coverslip slits with a double-sided tape as spacers. Self-organization of Min proteins inside the droplets was observed using the same equipment described for SLBs.

### Diffusion analysis

For analysis of diffusion of sfGFP in cytosolic parts and His-sfGFP-MinD on membranes in BSA solution entrapped inside microdroplets covered with *E. coli* polar lipids, a confocal laser-scanning microscope was used (FV1200; Olympus). The diffusion coefficient of sfGFP in 0 mg/mL, 50 mg/mL, 100 mg/mL, 200 mg/mL, and 300 mg/mL of BSA in RE buffer was measured by the standard protocol for Fluorescence Correlation Spectroscopy of FV1200. Diffusion coefficients of His-sfGFP-MinD on membranes in 0 mg/mL and 100 mg/mL of BSA in RE buffer was measured by Fluorescence Recovery After Photo-bleaching (FRAP) using tornado bleaching of circle area with ~1μm diameter. The recovery intensity as a function of time was converted to diffusion coefficients by using the FRAP protocol of FV1200.

### Pull-down assay

The mixture of 9 μM MinE-mCherry-His, 3 μM His-sfGFP-MinD, or 6 μM its mutant treated by thrombin (MinD^D40A^Δ10), and 3 μM BSA were applied to cOmplete His-Tag purification resin and incubated in RE buffer for 30 min at room temperature. Each mixture with resin was loaded into Micro Bio-Spin chromatography columns (Bio-Rad). Then, flow thorough fraction was separated and collected by a tabletop centrifuge. After washing the resin by 500 μL RE buffer with 20 mM imidazole for 3-5 times, elution fraction was obtained by 50 μL RE buffer with 250 mM imidazole. Proteins in each fraction were separated by SDS-PAGE and visualized by CBB staining.

### Evaluation of c/m ratio

Preparation of lipid droplets and glass coverslips for observation were performed using the same procedure as self-organization assay inside lipid droplets. To analyze localization (c/m) of MinE, various concentrations of MinE-mCherry-His (WT or I74M mutant) were mixed with BSA. For the analysis using *E. coli* cell extract, Ficoll70 (Santa Cruz Biotechnology, Dallas, TX, USA), or PEG8000 (Promega, Madison, WI, USA), 1 μM MinE-mCherry-His was used. Then, the mixture was entrapped inside microdroplets of *E. coli* polar lipids. To analyze localization of other proteins, 1 μM sfGFP-MinD with or without 50 mg/mL BSA, 1 μM sfGFP, or 1 μM BSA-FITC (Thermo Fisher Scientific, Waltham, MA, USA) with 1 mg/mL BSA were used. Membrane localization of each protein was observed using a confocal laser-scanning microscope FV1000 (Olympus). All analyses of obtained images were carried out using Fiji software. A center line of a droplet was manually drawn, and then, the intensity in each pixels of the line were obtained. The membrane intensities (m) were determined as the higher intensity of two membrane edge peaks. The cytosol intensities (c) were determined as the average intensity of 10 pixels around the pixel at the center position between two edge peaks. To evaluate the effects of macromolecules (Ficoll70, and PEG8000) and the modified lipid condition (85% of DOPC and 15% cardiolipin) in Figure 3F, c/m ratio was determined as the average from 10 individual droplets with 10-30 μm in a diameter.

### Numerical simulations

The partial differential equations in Model I (Eqs. (12)–(16)) and Model II (Eqs. (19)–(24)) were solved either using the commercial software of the Finite Element method, COMSOL, or using custom codes of the pseudo-spectral method in the spherical coordinates (*r*,*θ*,*φ*) for the closed membrane and Cartesian coordinates (*x*, *y*, *z*) for the planar membrane. Both methods reproduce waves in the planar membrane as well as waves in the closed membrane. In the pseudo-spectral method for the closed membrane of a spherical shape, all the concentration fields in bulks such as *c*_*D*_ and on a membrane such as *c*_*d*_ are expanded in terms of spherical harmonics, 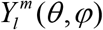:

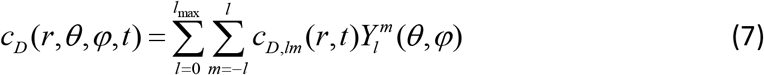

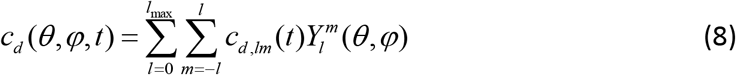

The model was then translated into a set of ordinary differential equations to obtain membrane concentrations (*c*_*d*_, *c*_*de*_, *c*_*e*_) and one-dimensional partial differential equations (time, *t*, and the radial direction, *r*) for the bulk concentrations (*c*_*D*_, *c*_*E*_). In total, we solved (*l*_max_ + 1)^2^ equations for each variable where the truncation of the mode was chosen as *l*_max_ = 16. The results were independent of the increase in the value.

Inhomogeneity of the concentration field on the membrane was expressed by the amplitude of each mode denoted by *l*. The amplitude is expressed by rotationally invariant form using the expansion coefficients in Eq.(7) with all *m*∊[−*l*,*l*]. For example, the uniform distribution of MinD on the membrane was expressed by the *l*=0 mode and its norm 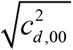, while the first mode (*l* = 1) corresponds to the inhomogeneous concentration field of a single wave, which is characterized by the norm 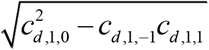.

The parameters were set as *ω*_*D*_ = 0.1, *ω*_*dD*_ = 5.0, *ω*_*E*_ = 0.1, *D* = 100, and *ω*_*ed*_ = 100 in the non-dimensional unit (see Appendix 2). If we choose *ω*_*e*_ = 0.2 [1/sec], *D*_*d*_ = 0.2 [μm^2^/sec], and the units of concentrations in cytosol and on membrane to be 10^3^ [1/μm^3^] and 10^3^ [1/μm^2^], respectively, then, our choice of the parameters implies *ω*_*D*_ = 0.02[μm/sec], *ω*_*dD*_ = 10^−3^ [μm^3^/sec], *ω*_*E*_ = 2 × 10^−5^ [μm^3^/sec], *D* = 20 [μm^2^/sec], and *ω*_*ed*_ = 2 × 10^−2^ [μm^2^/sec]. MinE localization at the membrane was modeled by the term *c*_*e*,0_. When BSA was added, we set smaller *c*_*e*,0_, whereas without BSA, we set *c*_*e*,0_ > 0. Note that when *c*_*e*,0_ = 0, all MinE molecules are in bulk *c*_*e*_ = 0 without MinD, while the membrane is filled by MinD, that is *c*_*d*_ = 1, without MinE.

For the planar membrane in the **x** = (*x*, *y*) plane, the concentration fields are expanded with the wave vector, **k**, such as

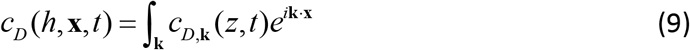

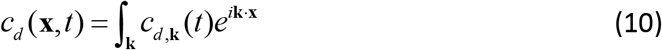

with the wave-number-dependent expansion coefficients *c*_*d,k*_(*t*) on the membrane and *c*_*D,k*_ (*z*, *t*) in the bulk cytosol. Here, the amplitude of the wave vector is denoted by the wave number, *k* = |**k**|. We may use the pseudo-spectral method, and solve the fields in the direction of the height, *z*, in real space, and the fields in the direction of the plane, (*x*, *y*), in Fourier space. The amplitude of a wave of MinD on the membrane is given by the absolute value of the complex number of the expansion coefficient |*c*_*d,k*_|.

#### Stability analysis of the theoretical models

We performed linear-stability analysis on the models. First, we calculated the stationary uniform solutions of the equations by setting time and spatial derivatives along the direction on the membrane to zero, and denoted these solutions by superscript “*”. Equations (12)–(16) and the boundary conditions [Eqs. (17) and (18)] were then linearized around the stationary uniform solution such as *c* = *c*\* *δc*. The eigenvalues, *σ*, are obtained by plugging *δc*(*t*) = *δce*^*σt*^ into the linearized equations (34, 45). The partial differential equations for the bulk dynamics were solved and the boundary conditions were translated into linear relationship between membrane and bulk concentrations. The set of the linearized equations for the concentration fields, for example **Ψ** = (*c*_*d*_, *c*_*de*_, *c*_*e*_, *c*_*D*_, *c*_*E*_) in Model I, is expressed by a matrix form as

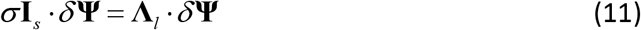

where the 5×5 matrix **Λ**_*l*_ has five eigenvalues depending on the mode *l* (but not on *m*) of spherical harmonics for the closed membrane. Here, **I**_*s*_ show the dynamics on the membrane and is a diagonal matrix whose diagonal elements are 1 only for the membrane concentrations and 0 otherwise, for example in Model I (1,1,1, 0, 0). The concentration in bulk in **Ψ** is interpreted as the concentration near the membrane, such that *c*_*D*_ (*R*,*θ*,*φ*) for the closed membrane and *c*_*D*_ (*x*, *y*, 0) for the planar membrane. For the planar membrane, the matrix is dependent on the wave number *k* and is denoted by **Λ**_*k*_. When the real part of the eigenvalue is positive, that is Re Λ_*l*_ > 0 for *l* ≠ 0, the uniform state is unstable, and an inhomogeneous pattern appears. Additionally, when the imaginary part is non-zero, the frequency becomes finite and either standing or rotating waves appear. In Model II, the same analysis was performed for the concentration fields denoted by **Ψ** = (*c*_*d*_, *c*_*de*_, *c*_*e*_, *c*_*DT*_ + *c*_*DD*_, *c*_*DD*_, *c*_*E*_) and the 6×6 matrix **Λ**_*l*_ in Eq.(11).

## Appendix

## Appendix 1 Theoretical analysis of the Min system of the closed and planar membranes

Spontaneous wave generation in the experiments have been studied as wave instability, which was proposed by Alan Turing as an extension of static Turing instability (46). In contrast with the static instability realized by a minimum of two components, wave instability requires three components. Although the instability can be evaluated by the linear stability analysis, physical intuition of the wave instability is not as obvious as static Turing instability. Therefore, several mechanisms have been proposed: First, homogeneous oscillation in a two-component system is suppressed by a third component (47), second, static inhomogeneous pattern generated by two-component Turing instability becomes oscillatory due to a third component (48) and, third, oscillatory instability occurs in eigenmodes associated with conserved quantities (49). Despite these phenomenology, clear mechanism of wave generation of Min system still remains under debate due to the complexity of reaction couplings and also lack of understanding how mixture of different spatial dimensions (membrane and cytosol) plays a role. The Min systems have two representative geometry; one is the closed membrane (Figure 7A), which we focused in this study, and the second is the open planar membrane (Figure 7B). In both systems, MinD and MinE proteins are distributed on the two-dimensional membrane and in the three-dimensional bulk cytosol.

So far, simple and realistic computational simulations have been proposed to reveal the mechanism of Min waves. These studies are based on partial differential equations of reaction-diffusion models (14, 30, 31, 34, 49–53) or particle-based stochastic models (54–57). The stochastic model is based on the model proposed by Huang et al. (31). At the early stage of the modeling, the geometry of the Min system was neglected such that all the concentration fields, both on membrane and in bulk cytosol, were defined in the same dimensions (50, 51). Recently, the coupling between the dynamics of membrane and bulk has been investigated. Among these models, only two approaches have successfully reported reproduction of the Min wave generation both on the planar and closed membrane including the effect of Min proteins in cytosol. One is to include transformation from ADP-MinD to ATP-MinD (31), and second is to include formation of a MinDE complex from membrane-bound MinD and MinE resulting in persistent MinE membrane binding (30). In both models, the Min wave occurs in the planar membrane (30, 34). The wave on the closed membrane requires a specific initial condition, stochasticity (54), or ellipsoidal shape (53) using the model proposed by (31), whereas the wave occurs without these effects in (30). The two approaches, however, show different dependence of the stability of the waves on total MinD and MinE concentrations, and other parameters.

## Appendix 2 Theoretical Model I

In Model I, MinD and MinE concentrations inside a spherical membrane with its radius *R*, or in the rectangular bulk with its height *H*, were denoted by *c*_D_ and *c*_E_, respectively (see Figure 7A and B). Concentrations of MinD, MinE, and their complex (MinDE) bound to a membrane were denoted by *c*_d_, *c*_e_, and *c*_de_, respectively (see Fig.S7A). The total MinD and MinE concentration is denoted by 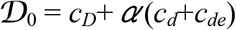 and 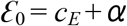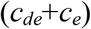, respectively. We denote the characteristic concentrations on the membrane and the cytosol as *c*_*s*_ and *c*_*b*_, respectively, and we express all the concentration fields in the unit of these characteristic concentrations. Here, *α* demonstrates an effect of confinement. Its concrete form is dependent on geometry of the system, but, in the current model for a spherical closed membrane, *α* = 3*c*_*s*_ / (*c*_*b*_*R*). For the planar membrane, it is associated with the height *H* of the system as *α* = *c*_*s*_ / (*c*_*b*_*H*).

Chemical reactions are schematically shown in Figure 5—figure supplement 1A. Each reaction shows a rate, *ω*, specified by its subscript. The diffusion constants of proteins bound to the membrane were denoted by *D*_*d*_, *D*_*e*_, and *D*_*de*_, whereas bulk diffusion of unbound proteins was denoted by the diffusion constants *D*_*D*_ and *D*_*E*_. We assumed the same diffusion constants for MinD and MinE in bulk represented by *D*. We also assumed the same diffusion constants for *D*_*d*_, *D*_*e*_, and *D*_*de*_ on the membrane. The latter diffusion constant was chosen to be unity without loss of generality. In comparison to the original work in (30), the unbinding process was approximated as *ω*_*de,m*_ = *ω*_*de*_ ≈ *ω*_e_, and *ω*_*de,c*_ = 0.

We defined the unit time scale as *τ*_0_ = 1/ *ω*_*e*_ and the unit length scale as 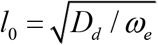. The concentration fields on the membrane were normalized by the characteristic concentration on the membrane, *c*_s_, which is chosen to be the maximum concentration on the membrane, *c*_max_, in the presence of the saturation effect (Model I). The model is given by the following equations (30):
Model I

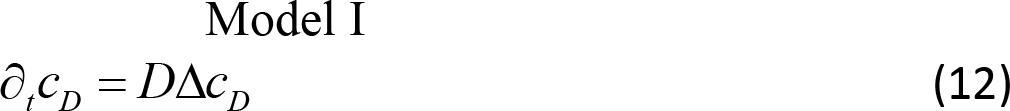

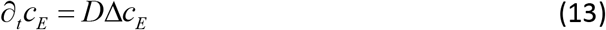

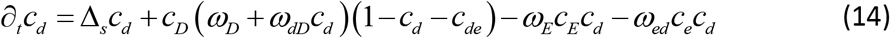

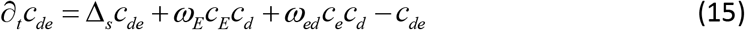

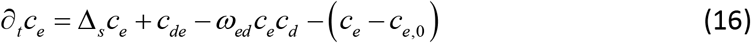

Here, Δ and Δ_s_ denote the Laplacian operator in three-dimensional bulk space and the Laplace-Bertrami operator on the two-dimensional surface, respectively. The boundary conditions of Eq.(12) and Eq.(13) are

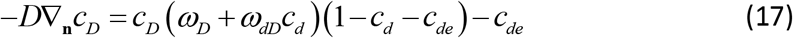

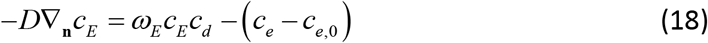

Here,∇_**n**_ is the derivative along the normal direction to the membrane. In this model, the set of concentration fields is expressed by Ψ = (*c*_*d*_, *c*_*de*_, *c*_*e*_, *c*_*D*_, *c*_*E*_) where the membrane concentration fields are *ψ* = (*c*_*d*_, *c*_*de*_, *c*_*e*_) and the bulk concentration fields are *ϕ* = (*c*_*D*_, *c*_*E*_).

## Appendix 3 Theoretical Model II

The effect of ATP hydrolization of MinD in bulk plays an essential role in the model proposed by Huang *et al.*(31). This model assumes MinE is in complex form of MinDE on the membrane. In a previous report(14), the model is generalized to include the effect of formation of MinDE complex from membrane-bound MinD and MinE. We, therefore, considered the following model:
Model II

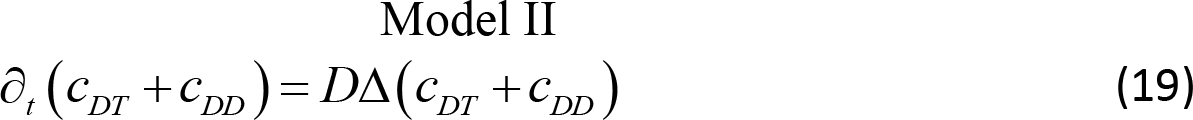

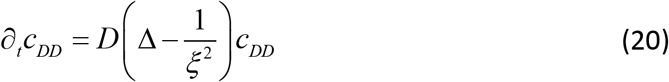

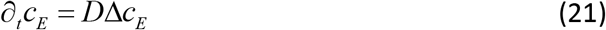

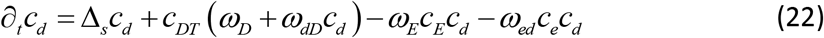

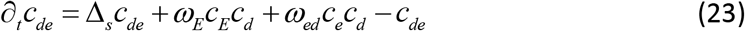

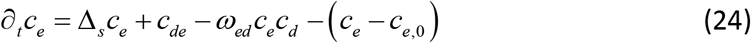

The length scale associated with ATP hydrolization is denoted by 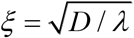, where *λ* is the rate of ATP hydrolization. The boundary conditions of Eqs.(19)-(21) are

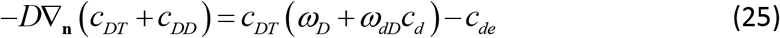

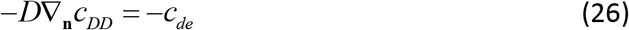

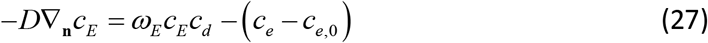

Chemical reactions are schematically shown in Figure 5—figure supplement 1B. Except ATP-hydrolization, this model differs from Eqs.(12)-(16) only in saturation of membrane-bound MinD in Eqs.(22) and (25). As we show in the analysis in Figure 6B, this effect is not relevant in closed membrane. In fact, when 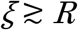, this model reproduces similar waves as in the model I, while when 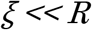 and *ω*_*ed*_ ≪1, this model reproduce a similar standing wave from the initial condition in which is *c*_*d*_ accumulated semi-sphere on the membrane (31, 54). We use the same parameters as Model I, and the additional parameter is set to be *λ* = 1.0. In this model, it is convenient to choose the set of concentration fields to be expressed by Ψ = (*c*_*d*_, *c*_*de*_, *c*_*e*_, *c*_*DT*_ + *c*_*DD*_, *c*_*DD*_, *c*_*E*_) where the membrane concentration fields are *ψ* = (*c*_*d*_, *c*_*de*_, *c*_*e*_) and the bulk concentration fields are *ϕ* = (*c*_*DT*_ + *c*_*DD*_, *c*_*DD*_, *c*_*E*_).

## Saturation of membrane-bound proteins does not play a role in the closed membrane

Model I differs from Model II in two respects. One is ATP hydrolysis in bulk proposed by (31). The Second effect is saturation membrane-bound MinD, that is, the concentration of MinD does not exceed a certain value (1 in our unit) which is given as a phenomenological parameter. This term was questioned by (58), in which ATP hydrolysis in bulk caps the concentration without this term. In order to show the saturation term is not necessary in a small system even without ATP hydrolysis in bulk, we compare stability analysis of Model I with and without the saturation term (Figure 6B and Figure 6—figure supplement 1). The results show they are almost identical, and the same mechanism of wave instability, namely suppression of instability at the zero mode, occurs in both cases. This is because maximum concentration of MinD on the membrane is not set by the saturation term rather by conservation law. For a larger system, this is not the case because the bulk concentrations is insensitive to the membrane concentrations due to small *α*.

## Appendix 4 Generic model and its reduction onto membrane

In order to give a unified expression for different model (Model I and II), we investigated a generic form of these models. We considered *n* variables of membrane-bound proteins denoted by *ψ* such as *ψ* = (*c*_*d*_, *c*_*de*_,···), and *m* variables of bulk cytosol concentrations denoted by *ϕ* such as *ϕ* = (*c*_*DD*_, *c*_*E*_,···). The dynamics in bulk is expressed by linear equations such as

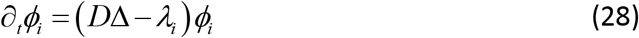

The subscript denoted a specific concentration of a protein in bulk cytosol. The dynamics of the membrane concentrations is formally written as

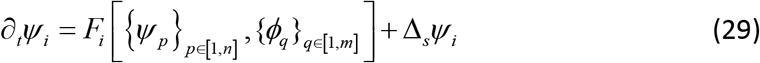

where the first term in the right-hand side expresses biochemical reactions. The boundary conditions are, in general, nonlinear, but they are rewritten as

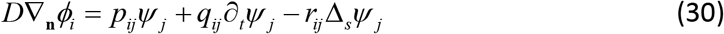

The matrices, **p**, **q**, and **r**, are specified by each model. Homogeneous stationary solution is obtain by *F*_*i*_[*ψ*\*,*ϕ*\*] = 0 together with conservation law of MinD and MinE.

The concentration fields in the linearized equation are expanded with spherical harmonics Eq.(8) for the closed membrane or with wave vectors **k** Eq.(10) for the planar membrane. Using the eigenvalues, the concentration fields are expressed as *ψ*(*t*) = *ψ*\* + *δψe*^*σt*^ on the membrane and *ϕ*(*t*) = *ϕ*\* + *δϕe*^*σt*^ in bulk. We can solve Eq.(28) together with the boundary conditions Eq.(30). Then the linearized equation on the membrane is expressed as

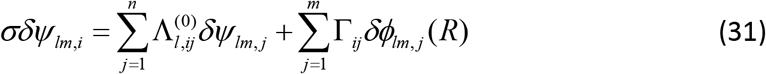

where the *n*×*n* matrix 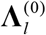 is expressed by

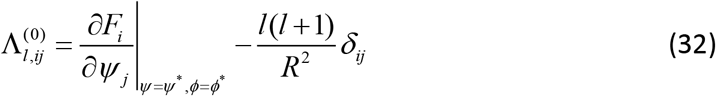

And the coupling between membrane and bulk dynamics is expressed by *n*×*m* matrix **Γ** as

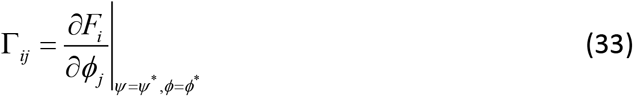

For the planar membrane, the wave-number-dependent concentrations *ϕ*_*k*_ and *ψ*_*k*_ were considered, and 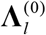 and *l*(*l* + 1) / *R*^2^ were replaced by 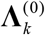 and *k*^2^, respectively. To obtain the eigenvalues, we solved the equation in which the determinant of the linear matrix in Eq.(31) vanished (see Eq.(1)). This is similar to Eq.(11), but it has a (*n* + *m*) × (*n* + *m*) matrix. The current form has only a *n*×*n* matrix, which makes the effect of confinement clearer as shown in Eq.(1).

Because there are two conserved quantities of this system, total MinD and MinE concentrations, there are two zero eigenvalues (zero eigenmodes) associated with them. Except these eigenvalues, the bottom-right block of the matrix **Λ**_*l*_ associated with bulk concentrations is invertible, and thus, we may eliminate the bulk concentrations by solving the linearized equations. This argument assumes that wave instability does not occur by the eigenvalues at the finite modes connected to the zero eigenmodes. This is the case in the models studied here, and the results in (34) for the planar membrane with the model proposed by (31) also demonstrate that the instability at a finite wavenumber is not connected to the zero eigenmodes.

## Supplementary Materials

Video 1. Behaviors of Min proteins entrapped in microdroplets

Video 2. Wave propagation of Min proteins in microdroplets with a lipid mixture (85% DOPC and 15% Cardiolipin)

Video 3. Wave propagation of Min proteins in microdroplets containing 100 mg/mL BSA

Video 4. Behaviors of Min proteins entrapped in microdroplets containing 100 mg/mL PEG8000

Video 5. Behaviors of Min proteins entrapped in microdroplets containing 100 mg/mL Ficoll70

Video 6. Wave propagation of non-tagged MinD tracked by sfGFP-MinC in microdroplets containing 100 mg/mL BSA

Video 7. Wave propagation of Min proteins in microdroplets containing 16 mg/mL macromolecules in cell extract

Video 8. Behaviors of sfGFP-MinD and MinEI74M-mCherry entrapped in microdroplets containing 50 mg/mL BSA

Video 9. Time development at initial stages of MinD single waves in lipids droplets using Model I (simulation)

Video 10. Time development at initial stages of MinD single waves in lipids droplets (experiment)

## Acknowledgments

We thank Ms. A. Yoshida (Keio University) for supporting protein purification, Prof. K. Yoshikawa (Doshisha University), Prof. H. Kitahata (Chiba University), Prof. T. Sakurai (Chiba University), Prof. Karsten Kruse (University of Geneva), and Prof. Toshiyuki Ogawa (Meiji University) for helpful discussion. We thank the financial support by JSPS KAKENHI Grant Number JP16H00809, JP26650044, JP15KT0081, JP15H00826, JP18H04565 for K.F., JP26800219, JP16H00793, and JP17K05605 for N.Y. We also thank Ph.D. Program Research Grant at Keio university for S.K.

---

**Figure 1—figure supplement 1:**
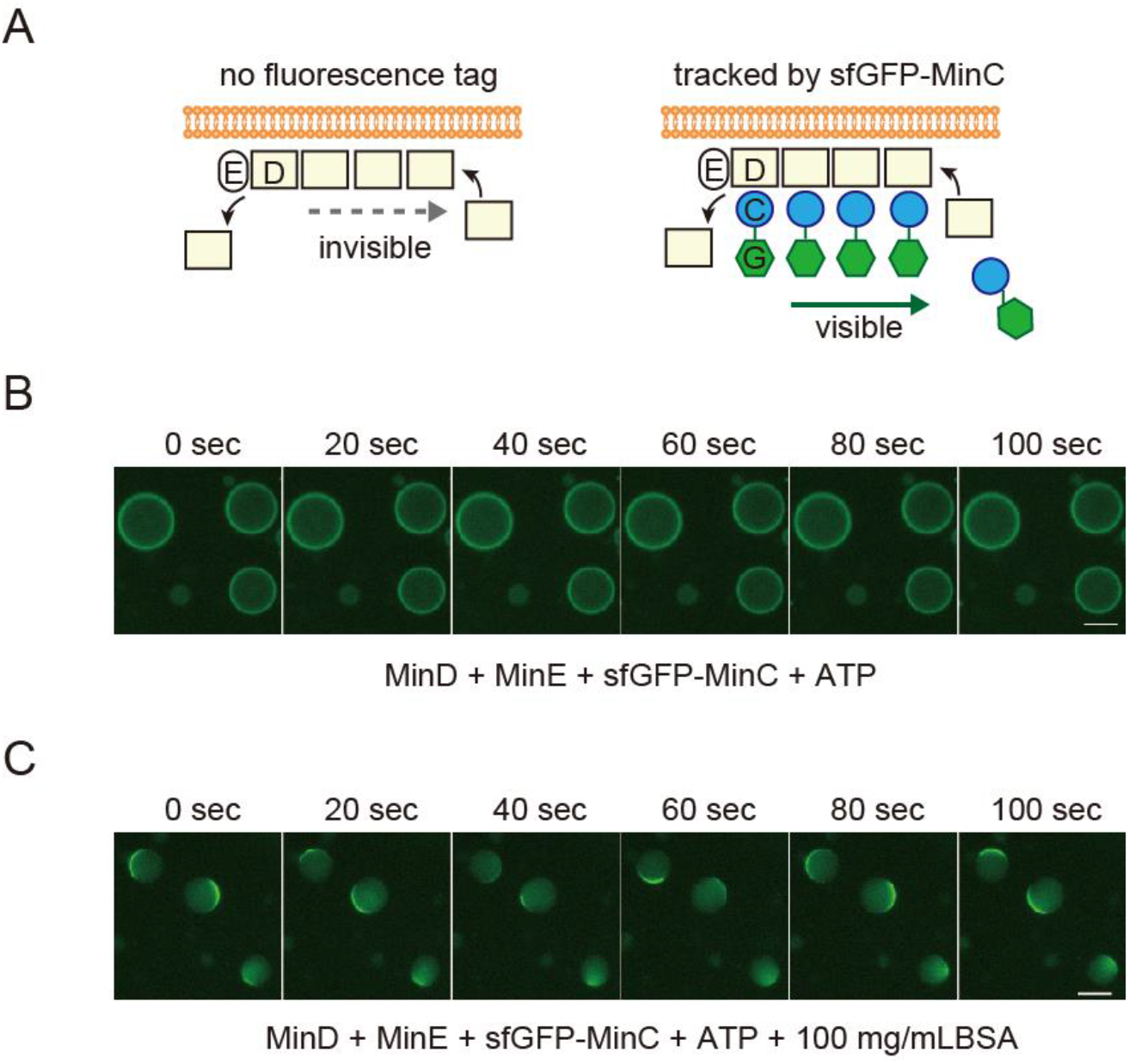
Tracking MinD by fluorescence-tagged MinC. (**A**) Representative illustration of MinD tracking by MinC fused with sfGFP at N-terminal. C, D, E, and G indicates MinC, MinD, MinE, and sfGFP. Because of interaction between MinD and MinC, sfGFP-MinC can track movement of no-tagged MinD. (**B**) and (**C**) indicates results of the time-lapse images of MinD tracked sfGFP-tracking without (**B**) or with (**C**) 100 mg/mL BSA.

**Figure 1—figure supplement 2:**
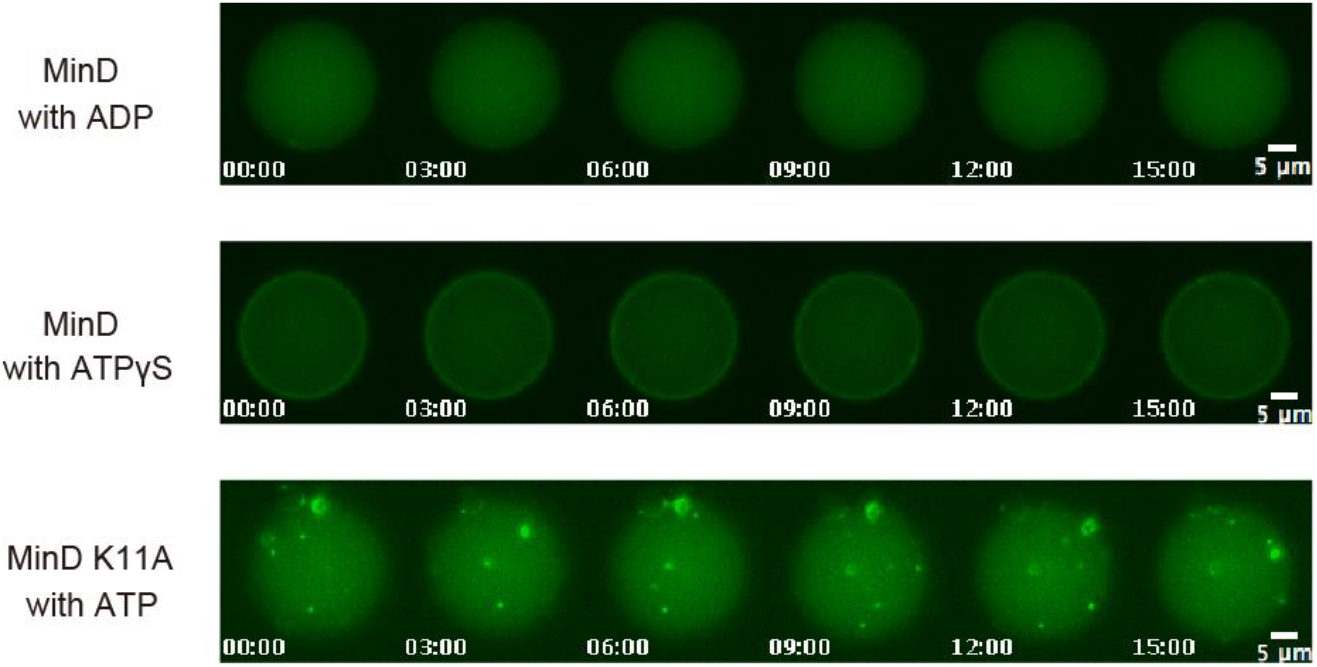
ATP dependence of the Min system on wave propagation in microdroplets containing 100 mg/mL BSA. ATP requirements of the Min wave were examined by replacing ATP with ADP or ATP_γ_S, and MinD with an ATPase-deficient MinD mutant (K11A) (28). Time-lapse images of MinD or its mutant in a representative droplet are shown.

**Figure 1—figure supplement 3:**
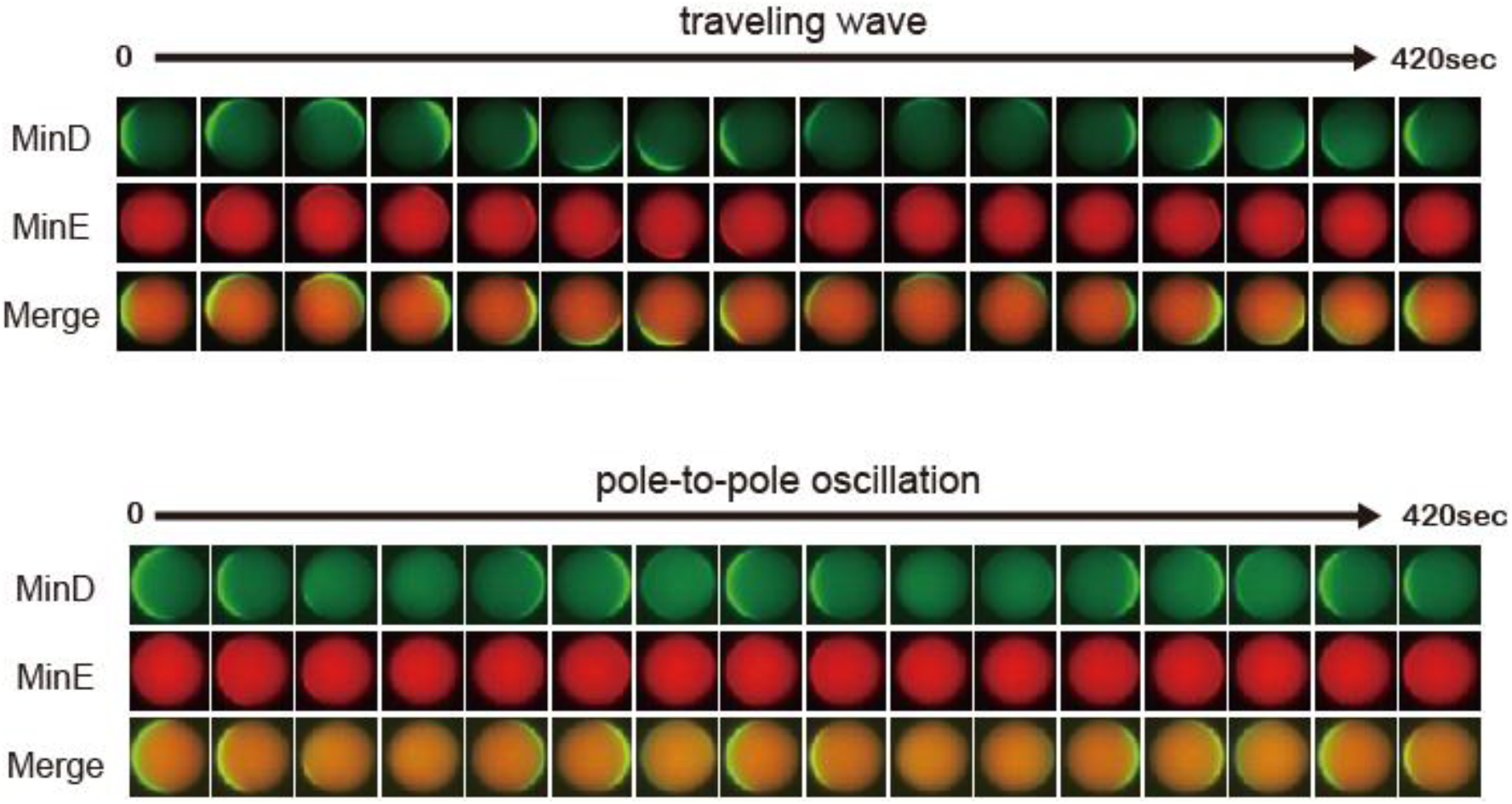
Time-lapse images of propagation waves in microdroplets. Representative time-lapse images of traveling wave and pole-to-pole oscillation in microdroplets were shown. Diameters of microdroplets are 25 μm (traveling waves) and 22 μm (pole-to-pole oscillation).

**Figure 2—figure supplement 1:**
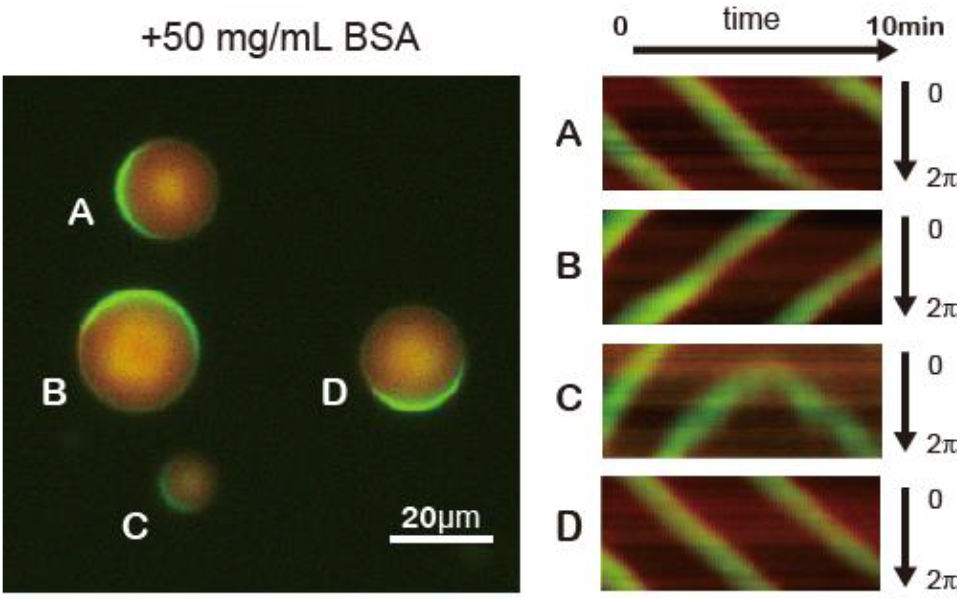
Traveling waves emerging in microdroplets containing 50 mg/mL BSA. Kymographs of MinD and MinE on the membrane of each droplet are shown. Green and red indicates sfGFP-MinD and MinE-mCherry, respectively.

**Figure 4—figure supplement 1:**
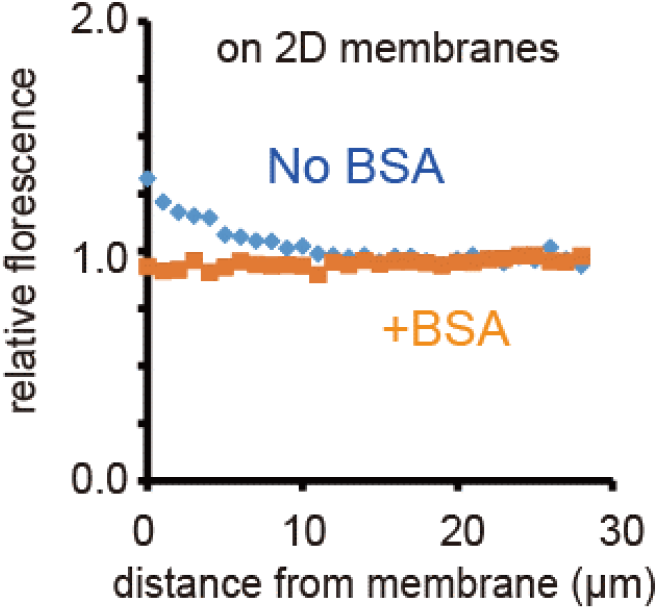
Localization of MinE near the two-dimensional planar membrane. Localization of MinE near the two-dimensional SLB was analysed by confocal microscopy. Relative fluorescence intensity plotted as a function of distance from the SLB. Fluorescence intensity was normalized by the value of 100 μm from two-dimensional SLB after subtraction of background noise.

**Figure 4—figure supplement 2:**
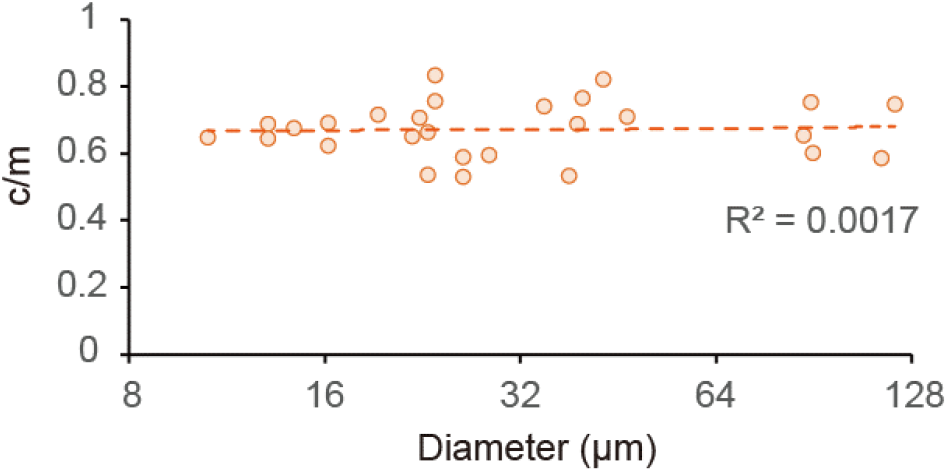
Size-dependence of c/m in microdroplets contacting 10 mg/mL BSA. Spontaneous localization rates (c/m) of MinE-mCherry in microdroplets contacting 10mg/mL BSA were plotted against sizes of microdroplets.

**Figure 5—figure supplement 1:**
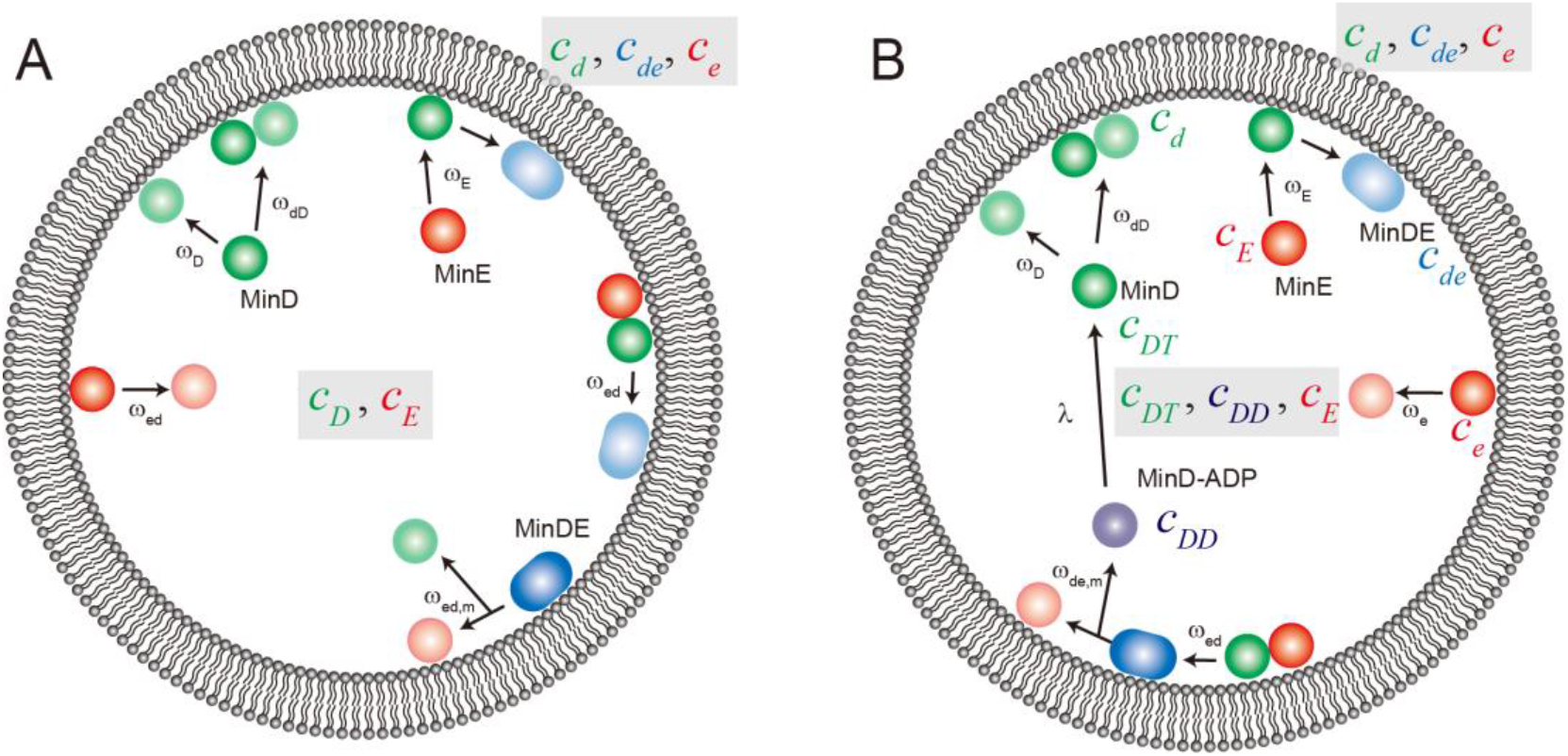
Reaction scheme for computational simulation of Min waves in cell-sized space. Schematic illustration of reaction constants used in our computational simulation model (**A** for model I, **B** for model II) are shown.

**Figure 6—figure supplement 1:**
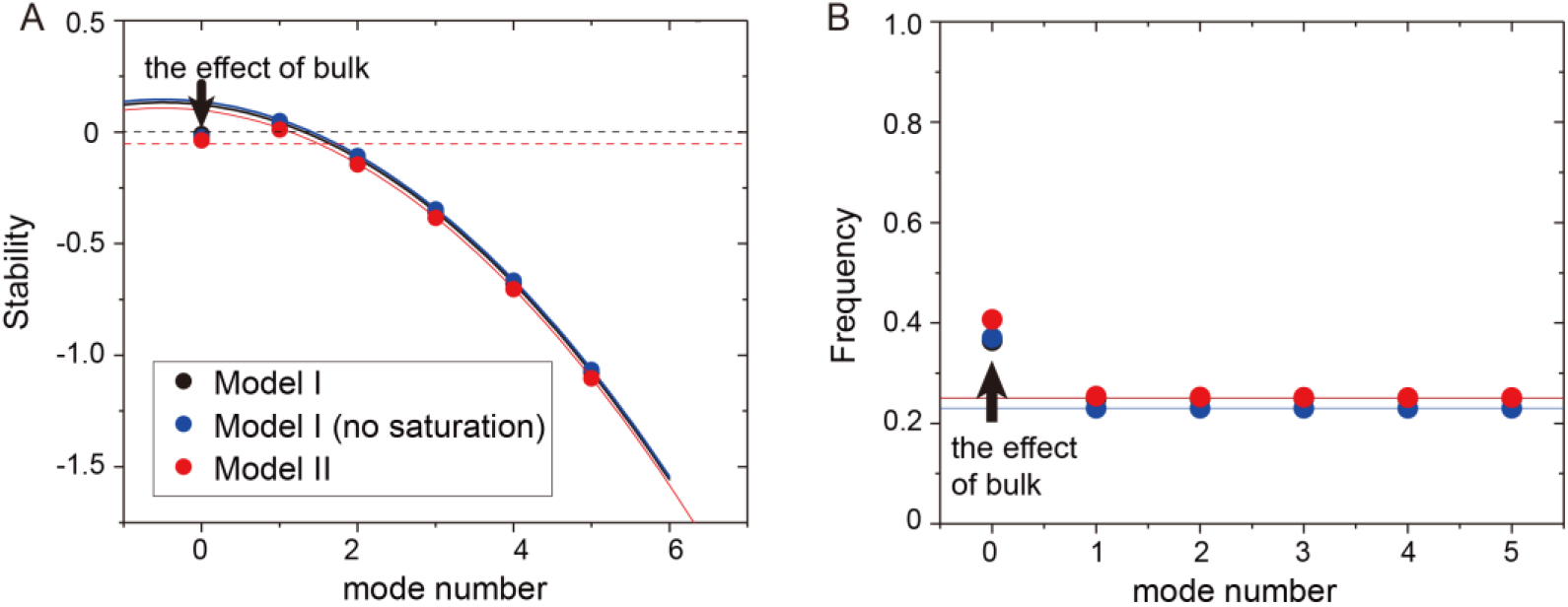
Mode dependence of stability and frequency of the different models obtained from linear stability analysis. (**A**) Real part of eigenvalues. The dashed black line denotes a zero eigenvalue. (**B**) imaginary part of eigenvalues. The stability and frequency obtained from the analysis without the effect of bulk is shown by solid lines for Model I (black), Model I without saturation term (blue), and Model II (red). The dashed red line in (**A**) shows the stability of the homogeneous state with the effect of confinement.

## References

1. Adachi S, Hori K, & Hiraga S (2006) Subcellular positioning of F plasmid mediated by dynamic localization of SopA and SopB. J Mol Biol 356(4):850–863.

2. Arai Y, et al. (2010) Self-organization of the phosphatidylinositol lipids signaling system for random cell migration. Proc Natl Acad Sci USA 107(27):12399–12404.

3. Huang CH, Tang M, Shi C, Iglesias PA, & Devreotes PN (2013) An excitable signal integrator couples to an idling cytoskeletal oscillator to drive cell migration. Nat Cell Biol 15(11):1307–1316.

4. Goryachev AB & Pokhilko AV (2008) Dynamics of Cdc42 network embodies a Turing-type mechanism of yeast cell polarity. FEBS Lett 582(10):1437–1443.

5. Rothfield L, Taghbalout A, & Shih YL (2005) Spatial control of bacterial division-site placement. Nat Rev Microbiol 3(12):959–968.

6. Rowlett VW & Margolin W (2013) The bacterial Min system. Curr Biol 23(13):R553–556.

7. Loose M, Fischer-Friedrich E, Ries J, Kruse K, & Schwille P (2008) Spatial regulators for bacterial cell division self-organize into surface waves in vitro. Science 320(5877):789–792.

8. Loose M, Fischer-Friedrich E, Herold C, Kruse K, & Schwille P (2011) Min protein patterns emerge from rapid rebinding and membrane interaction of MinE. Nat Struct Mol Biol 18(5):577–583.

9. Martos A, Petrasek Z, & Schwille P (2013) Propagation of MinCDE waves on free-standing membranes. Environ Microbiol 15(12):3319–3326.

10. Zieske K & Schwille P (2013) Reconstitution of pole-to-pole oscillations of min proteins in microengineered polydimethylsiloxane compartments. Angew Chem Int Ed 52(1):459–462.

11. Vecchiarelli AG, Li M, Mizuuchi M, & Mizuuchi K (2014) Differential affinities of MinD and MinE to anionic phospholipid influence Min patterning dynamics in vitro. Mol Microbiol 93(3):453–463.

12. Zieske K, Chwastek G, & Schwille P (2016) Protein patterns and oscillations on lipid monolayers and in microdroplets. Angew Chem Int Ed. 55(43):13455–13459

13. Caspi Y & Dekker C (2016) Mapping out Min protein patterns in fully confined fluidic chambers. Elife 5:e19271

14. Denk J, et al. (2018) MinE conformational switching confers robustness on self-organized Min protein patterns. Proc Natl Acad Sci USA 115(18):4553–4558.

15. Litschel T, Ramm B, Maas R, Heymann M, & Schwille P (2018) Beating Vesicles: Encapsulated Protein Oscillations Cause Dynamic Membrane Deformations. Angew Chem Int Ed 57(50):16286–16290.

16. Yanagisawa M, Sakaue T, & Yoshikawa K (2014) Characteristic behavior of crowding macromolecules confined in cell-sized droplets. Int Rev Cell Mol Biol 307:175–204.

17. Kuchler A, Yoshimoto M, Luginbuhl S, Mavelli F, & Walde P (2016) Enzymatic reactions in confined environments. Nat Nanotechnol 11(5):409–420.

18. Watanabe C & Yanagisawa M (2018) Cell-size confinement effect on protein diffusion in crowded poly(ethylene)glycol solution. Phys Chem Chem Phys 20(13):8842–8847.

19. Zhabotinsky AM, Dolnik M, & Epstein IR (1995) Pattern formation arising from wave instability in a simple reaction—diffusion system. J Chem Phys 103(23):10306–10314.

20. Epstein IR & Showalter K (1996) Nonlinear chemical dynamics: Oscillations, patterns, and chaos. J Phys Chem 100(31):13132–13147.

21. Lee KJ, Mccormick WD, Pearson JE, & Swinney HL (1994) Experimental-observation of self-replicating spots in a reaction-diffusion system. Nature 369(6477):215–218.

22. Fujiwara K & Yanagisawa M (2014) Generation of giant unilamellar liposomes containing biomacromolecules at physiological intracellular concentrations using hypertonic conditions. ACS Synth Biol 3(12):870–874.

23. Zhou HX, Rivas G, & Minton AP (2008) Macromolecular crowding and confinement: biochemical, biophysical, and potential physiological consequences. Annu Rev Biophys 37:375–397.

24. Groen J, et al. (2015) Associative interactions in crowded solutions of biopolymers counteract depletion effects. J Am Chem Soc 137(40):13041–13048.

25. Schweizer J, et al. (2012) Geometry sensing by self-organized protein patterns. Proc Natl Acad Sci USA 109(38):15283–15288.

26. Martos A, et al. (2015) FtsZ polymers tethered to the, embrane by ZipA are susceptible to spatial regulation by Min waves. Biophys J 108(9):2371–2383.

27. Shih YL, Fu X, King GF, Le T, & Rothfield L (2002) Division site placement in *E.coli*: mutations that prevent formation of the MinE ring lead to loss of the normal midcell arrest of growth of polar MinD membrane domains. EMBO J 21(13):3347–3357.

28. Zhou H, et al. (2005) Analysis of MinD mutations reveals residues required for MinE stimulation of the MinD ATPase and residues required for MinC interaction. J Bacteriol 187(2):629–638.

29. Park KT, Villar MT, Artigues A, & Lutkenhaus J (2017) MinE conformational dynamics regulate membrane binding, MinD interaction, and Min oscillation. Proc Natl Acad Sci USA 114(29):7497–7504.

30. Bonny M, Fischer-Friedrich E, Loose M, Schwille P, & Kruse K (2013) Membrane binding of MinE allows for a comprehensive description of Min-protein pattern formation. PLoS Comput Biol 9(12):e1003347.

31. Huang KC, Meir Y, & Wingreen NS (2003) Dynamic structures in *Escherichia coli*: spontaneous formation of MinE rings and MinD polar zones. Proc Natl Acad Sci USA 100(22):12724–12728.

32. Hsieh CW, et al. (2010) Direct MinE-membrane interaction contributes to the proper localization of MinDE in *E. coli*. Mol Microbiol 75(2):499–512.

33. Vecchiarelli AG, Li M, Mizuuchi M, Ivanov V, & Mizuuchi K (2017) MinE recruits, stabilizes, releases, and inhibits MinD interactions with membrane to drive oscillation. bioRxiv:109637.

34. Halatek J & Frey E (2018) Rethinking pattern formation in reaction–diffusion systems. Nat Phys 14(5):507.

35. Pismen LM (2006) Patterns and interfaces in dissipative dynamics (Springer Science & Business Media).

36. Keener JP (1980) Waves in excitable media. SIAM J Appl Math 39(3):528–548.

37. Bär M & Eiswirth M (1993) Turbulence due to spiral breakup in a continuous excitable medium. Phys Rev E 48(3):R1635.

38. Winfree AT (1991) Varieties of spiral wave behavior: An experimentalist’s approach to the theory of excitable media. Chaos 1(3):303–334.

39. Ruggeri F, et al. (2013) Non-specific interactions between soluble proteins and lipids induce irreversible changes in the properties of lipid bilayers. Soft Matter 9(16):4219–4226.

40. Miyazaki M, Chiba M, Eguchi H, Ohki T, & Ishiwata S (2015) Cell-sized spherical confinement induces the spontaneous formation of contractile actomyosin rings in vitro. Nat Cell Biol 17(4):480–489.

41. Yanagisawa M, Nigorikawa S, Sakaue T, Fujiwara K, & Tokita M (2014) Multiple patterns of polymer gels in microspheres due to the interplay among phase separation, wetting, and gelation. Proc Natl Acad Sci USA 111(45):15894–15899.

42. Good MC, Vahey MD, Skandarajah A, Fletcher DA, & Heald R (2013) Cytoplasmic volume modulates spindle size during embryogenesis. Science 342(6160):856–860.

43. Fujiwara K & Doi N (2016) Biochemical Preparation of Cell Extract for Cell-Free Protein Synthesis without Physical Disruption. PLoS One 11(4):e0154614.

44. Fujiwara K & Nomura SM (2013) Condensation of an additive-free cell extract to mimic the conditions of live cells. PLoS One 8(1):e54155.

45. Gou J, Li Y, Nagata W, & Ward M (2015) Synchronized oscillatory dynamics for a 1-D model of membrane kinetics coupled by linear bulk diffusion. SIAM J Appl Dyn Syst 14(4):2096–2137.

46. Turing AM (1952) The chemical basis of morphogenesis. Phil Trans R Soc Lond B 237(641):37–72.

47. Gourley S & Britton N (1996) A predator-prey reaction-diffusion system with nonlocal effects. J Math Biol 34(3):297–333.

48. Okuzono T & Ohta T (2003) Traveling waves in phase-separating reactive mixtures. Phys Rev E 67(5 Pt 2):056211.

49. Kessler DA & Levine H (2016) Nonlinear self-adapting wave patterns. New Journal of Physics 18(12):122001.

50. Meinhardt H & de Boer PA (2001) Pattern formation in *Escherichia coli*: a model for the pole-to-pole oscillations of Min proteins and the localization of the division site. Proc Natl Acad Sci USA 98(25):14202–14207.

51. Howard M, Rutenberg AD, & de Vet S (2001) Dynamic compartmentalization of bacteria: accurate division in *E. coli*. Phys Rev Lett 87(27 Pt 1):278102.

52. Huang KC & Wingreen NS (2004) Min-protein oscillations in round bacteria. Phys Biol 1(3-4):229–235.

53. Halatek J & Frey E (2012) Highly canalized MinD transfer and MinE sequestration explain the origin of robust MinCDE-protein dynamics. Cell Rep 1(6):741–752.

54. Fange D & Elf J (2006) Noise-induced Min phenotypes in *E. coli*. PLoS Comput Biol 2(6):e80.

55. Kerr RA, Levine H, Sejnowski TJ, & Rappel WJ (2006) Division accuracy in a stochastic model of Min oscillations in *Escherichia coli*. Proc Natl Acad Sci USA 103(2):347–352.

56. Arjunan SN & Tomita M (2010) A new multicompartmental reaction-diffusion modeling method links transient membrane attachment of *E. coli* MinE to E-ring formation. Syst Synth Biol 4(1):35–53.

57. Hoffmann M & Schwarz US (2014) Oscillations of Min-proteins in micropatterned environments: a three-dimensional particle-based stochastic simulation approach. Soft Matter 10(14):2388–2396.

58. Halatek J & Frey E (2014) Effective 2D model does not account for geometry sensing by self-organized proteins patterns. Proc Natl Acad Sci USA 111(18):E1817.

